# Hyperactive MEK1 signaling in cortical GABAergic neurons causes embryonic parvalbumin-neuron death and defects in behavioral inhibition

**DOI:** 10.1101/748087

**Authors:** Michael C. Holter, Lauren T. Hewitt, Kenji J. Nishimura, George R. Bjorklund, Shiv Shah, Noah R. Fry, Katherina P. Rees, Tanya A. Gupta, Carter W. Daniels, Guohui Li, Steven Marsh, David M. Treiman, M. Foster Olive, Trent R. Anderson, Federico Sanabria, William D. Snider, Jason M. Newbern

**Affiliations:** School of Life Sciences, Arizona State University; Tempe, AZ 85287, USA; Department of Psychology, Arizona State University; Tempe, AZ 85287, USA; College of Medicine, University of Arizona; Phoenix, AZ 85004, USA; Barrow Neurological Institute; Phoenix, AZ 85013, USA; University of North Carolina Neuroscience Center, The University of North Carolina School of Medicine; Chapel Hill, NC, 27599, USA

**Author notes:** Interdepartmental Neuroscience Graduate Program, University of Texas; Austin, TX, 78712, USA. Department of Psychiatry, Columbia University, New York, NY 10032, USA. Corresponding Author address: School of Life Sciences, PO Box 84501 Arizona State University Tempe, AZ 85287-4501, USA.

**Keywords:** ERK1/2, cerebral cortex, development, ganglionic eminence, ADHD, RASopathy, kinase, apoptosis, response inhibition capacity

## Abstract

Abnormal ERK/MAPK pathway activity is an important contributor to the neuropathogenesis of many disorders including Fragile X, Rett, 16p11.2 Syndromes, and the RASopathies. Individuals with these syndromes often present with intellectual disability, ADHD, autism, and epilepsy. However, the pathological mechanisms that underly these deficits are not fully understood. Here, we examined whether hyperactivation of MEK1 signaling modifies the development of GABAergic cortical interneurons (CINs), a heterogeneous population of inhibitory neurons necessary for cortical function. We show that GABAergic-neuron specific MEK1 hyperactivation *in vivo* leads to increased cleaved caspase-3 labeling in a subpopulation of immature neurons in the embryonic subpallium. Adult mutants displayed a significant loss of mature parvalbumin-expressing (PV) CINs, but not somatostatin-expressing CINs, during postnatal development and a modest reduction in perisomatic inhibitory synapse formation on excitatory neurons. Surviving mutant PV-CINs maintained a typical fast-spiking phenotype and minor differences in intrinsic electrophysiological properties. These changes coincided with an increased risk of seizure-like phenotypes. In contrast to other mouse models of PV-CIN loss, we discovered a robust increase in the accumulation of perineuronal nets, an extracellular structure thought to restrict plasticity in the developing brain. Indeed, we found that mutants exhibit a significant impairment in the acquisition of a behavioral test that relies on behavioral response inhibition, a process linked to ADHD-like phenotypes. Overall, our data suggests PV-CIN development is particularly sensitive to hyperactive MEK1 signaling which may underlie neurological deficits frequently observed in ERK/MAPK-linked syndromes.

**Significance Statement:** The RASopathies are a family of neurodevelopmental syndromes caused by mutations that lead to increased RAS/RAF/MEK/ERK signaling and are associated with intellectual disability, epilepsy, and ADHD. We do not fully understand how distinct neuronal subtypes are affected in these syndromes. Here, we show that increased MEK signaling in developing mice promotes the embryonic death of a specific subset of cortical inhibitory neurons that express parvalbumin. Surviving mutant parvalbumin neurons also show significant changes in crucial maturation processes, which coincide with increased seizure susceptibility and profound deficits in behavioral inhibition. These data suggest that deficits in inhibitory circuit development contribute to RASopathy neuropathogenesis and indicate that therapeutic strategies targeting inhibitory interneuron dysfunction may be beneficial for these individuals.

## Introduction

Multiple developmental disorders are caused by genetic mutations linked to perturbation of kinase activity and altered intracellular signaling. The RAS/RAF/MEK/ERK (ERK/MAPK) pathway is a well-known, ubiquitous signaling cascade that is dynamically activated during development (Krens et al., 2006; Samuels et al., 2009). Mutations in classic RAS/MAPK signaling pathway components or upstream regulators, such as *PTPN11/SHP2*, *NF1*, or *SYNGAP1*, cause a family of related syndromes, known collectively as the RASopathies (Tidyman and Rauen, 2016). Moreover, *MAPK3/ERK1* is present in a frequently mutated region of 16p11.2 linked to Autism Spectrum Disorder (ASD) and animal models of Fragile X, Rett, and Angelmann Syndromes also exhibit changes in ERK/MAPK signaling activity (Kumar et al., 2008; Pucilowska et al., 2015; Vorstman et al., 2006). These disorders are often associated with intellectual disability, neurodevelopmental delay, ADHD, autism, and epilepsy. Clearly, aberrant ERK/MAPK activity is an important molecular mediator of neurodevelopmental abnormalities, however, therapeutic approaches for these conditions are lacking, due in part to a limited understanding of the developmental stage-and cell-specific functions of ERK/MAPK signaling in the brain. Delineating the precise consequences of altered ERK/MAPK activity on specific neuronal subtypes in the developing forebrain may provide insight into the neuropathogenesis of multiple neurodevelopmental diseases.

Coordinated interactions between multiple cell types are necessary for normal brain function, but deficits in select cellular subtypes often mediate specific neurodevelopmental phenotypes. Past work has shown that ERK/MAPK signaling regulates the development of dorsal cortex-derived glutamatergic cortical projection neurons (PNs) and glia (Aoidi et al., 2018; Ehrman et al., 2014; Ishii et al., 2013; Li et al., 2012). Upstream regulators, such as *Syngap1*, are also crucial for the early development of cortical glutamatergic neuron structure, excitability, and cognition (Clement et al., 2012; Ozkan et al., 2014). In contrast, NF1 mutations have been shown to impair aspects of spatial learning and memory via disruption of GABAergic, but not glutamatergic, neuron function (Cui et al 2008). Abnormal GABAergic circuitry is thought to be a key feature in the neuropathogenesis of various other neurodevelopmental disorders (Chao et al., 2010; Cui et al., 2008; Paluszkiewicz et al., 2011; Zhang et al., 2010). GABAergic neuron-directed *Syngap1* loss modulates GABAergic output but does not drive major abnormalities in mouse behavior or seizure threshold (Berryer et al., 2016; Ozkan et al., 2014). Mutations in signaling components upstream of *Ras* differentially modulate multiple downstream pathways and do not provide a clear delineation of ERK/MAPK function (Anastasaki and Gutmann, 2014; Brown et al., 2012). Here, we have hyperactivated MEK1 specifically in GABAergic cortical interneurons (CINs) to better understand the effects on inhibitory circuit development.

In the mature cortex, locally connected parvalbumin-(PV) and somatostatin-expressing (SST) CINs comprise the most populous and functionally diverse GABAergic subtypes (Kessaris et al., 2014). Reduced PV-CIN number is often observed in mouse models of multiple neurodevelopmental diseases, however, the mechanism of loss is poorly understood (Chao et al., 2010; Cui et al., 2008; Steullet et al., 2017). PV-and SST-CINs are generated in spatiotemporal fashion primarily from the medial ganglionic eminence (MGE) and migrate tangentially to the cortical plate (Gelman et al., 2009; Gelman and Marín, 2010; Lavdas et al., 1999; Marin and Rubenstein, 2001, 2003; Parnavelas, 2000; Tamamaki et al., 1997; Wichterle et al., 1999; Wichterle et al., 2001; Wonders and Anderson, 2006). Tangential migration and early GABAergic circuit development is regulated by BDNF/TRKB, GDNF/GFRα1, HGF/MET, and NRG/ERBB4 signaling, which activate multiple Receptor Tyrosine Kinase (RTK)-linked intracellular kinase cascades, including ERK/MAPK (Bae et al., 2010; Fazzari et al., 2010; Flames et al., 2004; Perrinjaquet et al., 2011; Pozas and Ibanez, 2005). While the transcriptional basis of GABAergic neuron development has been well-studied (Lim et al., 2018; Mayer et al., 2018; Mi et al., 2018; Paul et al., 2017), the kinase cascades that mediate GABAergic development in response to critical extracellular cues have received less attention.

Here, we show that GABAergic neuron-specific expression of constitutively-active MEK1^S217/221E^ (caMEK1) led to caspase-3 activation in a subset of embryonic GABAergic neurons. Even though caMEK1 is expressed in all CINs, we only observed a significant reduction in the number of mature PV-CINs, but not SST-CINs. In contrast with past models exhibiting PV-CIN loss, we found a surprising increase in the extent of perineuronal net (PNN) accumulation around these cells (Steullet et al., 2017). We observed an increased risk of spontaneous epileptiform activity and mild seizure-like activity in a subset of mutant mice that coincided with a reduction in inhibitory synapses on pyramidal neurons. Mutant mice exhibited normal locomotor, anxiety-like, and social behaviors, but we discovered deficits in behavioral response inhibition capacity, a process linked to ADHD-like phenotypes. Our findings indicate that GABAergic-specific MEK1 hyperactivation is sufficient to drive widespread changes in cortical development relevant to cognitive phenotypes observed in RASopathies. Together, these data define the precise functions of ERK/MAPK signaling in CIN development and suggest preferential contributions of PV-CIN pathology to ERK/MAPK-linked disorders.

## Results

### Differential expression of ERK/MAPK components in CINs

The ERK/MAPK cascade is a commonly utilized intracellular signaling pathway that is dynamically activated during embryogenesis and in adulthood. In the embryonic ventricular zone, neural progenitors typically show high levels of P-ERK1/2 relative to immature post-mitotic neurons (Pucilowska et al., 2018; Stanco et al., 2014). In adult cortices, elevated P-ERK1/2 labeling is enriched in a heterogeneous set of excitatory PNs, primarily in layer 2, and plays a critical role in long-range PN development (Cancedda et al., 2003; Gauthier et al., 2007; Holter et al., 2019; Pham et al., 2004; Pucilowska et al., 2012; Suzuki et al., 2004; Xing et al., 2016). The activation of ERK1/2 in CINs has not been well characterized. We generated mice expressing *Slc32A1:Cre* and the Cre-dependent red fluorescent protein (RFP) reporter, *Ai9* (Madisen et al., 2010; Vong et al., 2011) (Figure1A-D). As expected, brain regions abundant with GABA-expressing neurons robustly expressed *Ai9* (Figure 1A). Immunolabeling for MAP2K1 (MEK1) revealed relatively lower expression in CINs in comparison to NEUN^+^/RFP^-^ presumptive PNs (Figure 1E-G). Layer II/III CINs also expressed low levels of MAPK1/ERK2 in comparison to PNs (Figure 1H-J). In agreement with previous studies, high levels of P-ERK1/2 were observed in a subset of PNs in cortical layer II/III (Cancedda et al., 2003; Pham et al., 2004; Suzuki et al., 2004). However, examination of P-ERK1/2 immunolabeling in RFP^+^ CINs revealed qualitatively lower levels of P-ERK1/2 in comparison to PNs (Figure 1K-M). In summary, CINs express relatively lower levels of MEK1, ERK2, and P-ERK1/2 than excitatory neurons in the adult cortex, raising the possibility of functionally distinct roles for this cascade between these two primary cortical neuron subtypes.

**Figure 1.**
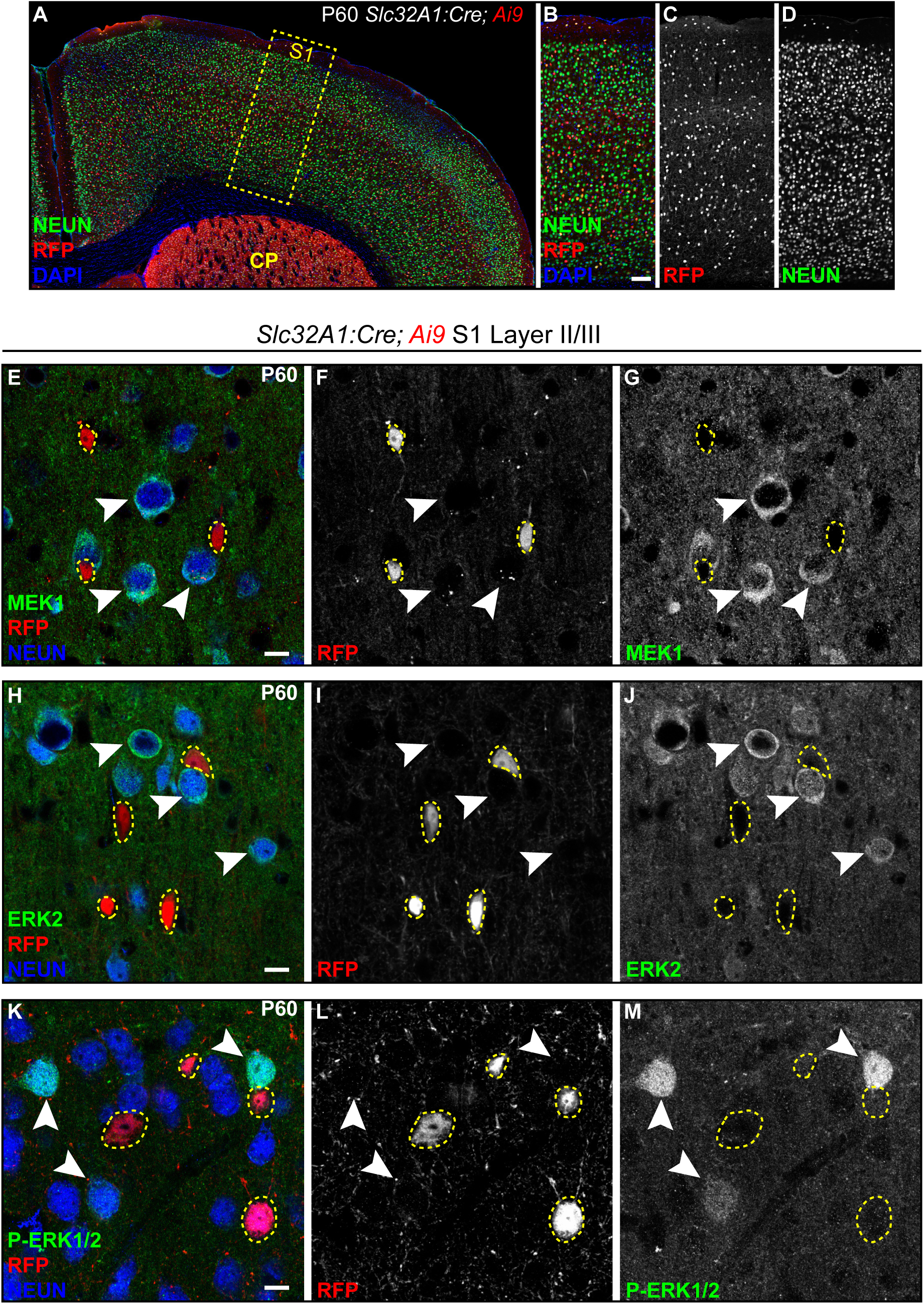
Cortical CINs exhibit low levels of ERK/MAPK expression and activity. (**A-D**) Representative confocal images of *Slc32A1:Cre, Ai9* sensorimotor cortex. Note the robust expression of RFP in brain regions with high densities of GABAergic neurons. (Scale bar = 100 µm) (**E-M**) Immunolabeling for MEK1 (E-G) and ERK2 (H-J) showed comparatively low expression in inhibitory CINs (yellow outlines) when compared to NEUN^+^/*Ai9*^-^ excitatory neurons (arrowheads) in layer II (n=3). Relatively lower expression of P-ERK1/2 was also detected in inhibitory CINs when compared to excitatory neurons (n=3). (Scale bar = 10 µm).

### GABAergic-autonomous caMEK1 expression decreases PV-CIN number

Increased ERK/MAPK signaling is the most common result of RASopathy-linked mutations (Tidyman and Rauen, 2016). We utilized a Cre-dependent, constitutively active *CAG-Loxp-Stop-Loxp-Mek1^S217/221E^* (*caMek1*) allele, which has been shown to hyperactivate MEK1/2-ERK1/2 signaling (Alessi et al., 1994; Bueno et al., 2000; Cowley et al., 1994; Klesse et al., 1999; Krenz et al., 2008; Lajiness et al., 2014; Li et al., 2012). We generated *caMek1*, *Slc32A1:Cre* mice to hyperactivate MEK1 in a CIN-specific fashion during embryogenesis. Elevated MEK1 expression was clearly detectable in the E13.5 mantle zone of the ganglionic eminences, presumptive embryonic CINs migrating into the cortex, and adult CINs in primary somatosensory cortex (S1) (Figure S1A-H; J-O). *CaMek1*, *Slc32A1:Cre* mice were viable and phenotypically normal, though mutants exhibited larger body mass than controls in adulthood (Figure S1I).

Surprisingly, assessment of fluorescently-labeled CINs in *caMek1*, *Slc32A1:Cre*, *Ai9* sensory cortices revealed a significant reduction in total RFP^+^ cell density (Figure 2A-I). In the adult cortex, approximately 40% of CINs express PV whereas 30% express SST, which serve as mostly non-overlapping markers of two distinct populations of CINs (Kelsom and Lu, 2013; Kessaris et al., 2014; Rudy et al., 2011). Strikingly, we observed a significant reduction in the proportion of PV^+^/RFP^+^ CINs, but not in the proportion of SST^+^/RFP^+^ CINs (Figure 2K-Q). PV, but not SST, expressing CINs displayed a significant increase in somal area compared to control neurons (Figure 2R-U, V). A reduced density of PV-CINs was detected in a separate *caMek1, Dlx5/6:Cre* strain that also targets postmitotic CINs in the developing cortex (Figure S2A-D) (Monory et al., 2006).

**Figure 2.**
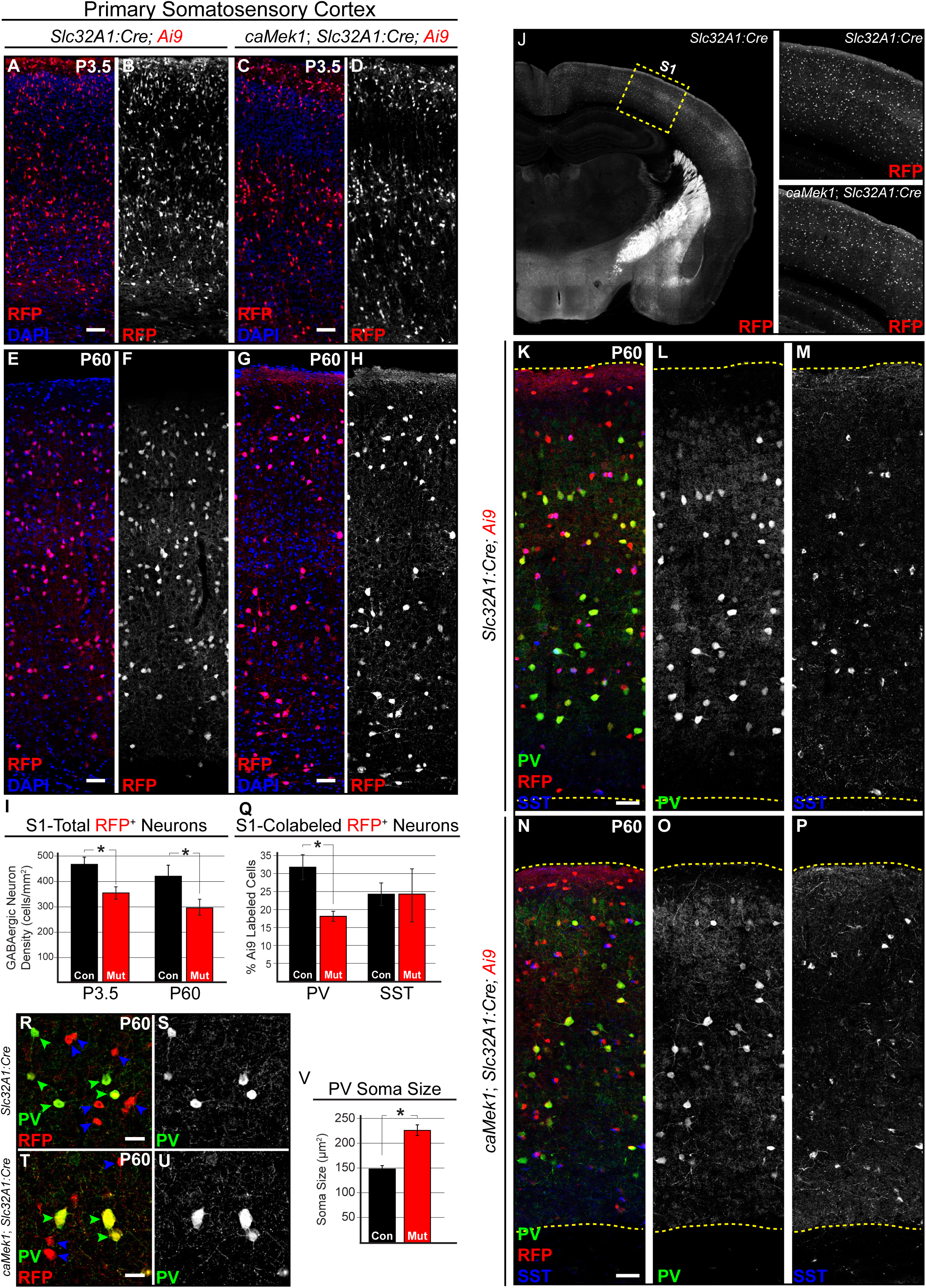
MEK1 hyperactivation leads to a selective reduction in PV-expressing CINs in the postnatal cortex. **(A-H)** *caMek1*, *Slc32A1:Cre*, *Ai9* mutant P3.5 (A-D) and P60 (E-H) primary somatosensory cortices exhibit reduced numbers of *Ai9*-expressing CINs in comparison to *Slc32A1:Cre*, *Ai9* controls (quantification in **I**, n=3, mean ± SEM, * = p < 0.05). (Scale bar = 100 µm) **(J-Q)** We quantified the proportion of fluorescently co-labeled PV/RFP or SST/RFP co-expressing CINs in the sensory cortex (J). Confocal micrographs of RFP-expressing CINs at P60 demonstrates that the proportion of PV^+^/RFP^+^ CINs, but not SST^+^/RFP^+^ CINs, was significantly decreased in mutants (N-P) in comparison to controls (K-M) (quantification in **Q**: n=3, mean ± SEM, * = p < 0.05). (Scale bar = 100 µm) **(R-V)** Mutant PV-CINs (T-U, green arrowheads in T) display increased soma size in comparison to control PV-CINs (R-S) (quantification in **V**, n = 21 control neurons, 42 mutant neurons, mean ± SEM, * = p < 0.001). PV^-^/RFP^+^ CINs displayed no qualitative change in soma size (blue arrowheads). (Scale bar = 25 µm).

Recombination within the entire GABAergic system left open the possibility that indirect changes in other neuroanatomical regions could alter global cortical activity and modulate PV-CIN number (Denaxa et al., 2018). To restrict Cre expression to primarily MGE-derived CINs, we generated *caMek1*, *Nkx2.1:Cre, Ai9* mice and assessed the proportion of PV^+^/RFP^+^ CINs. *CaMek1*, *Nkx2.1:Cre* mice exhibited generalized growth delay in the second-third postnatal week and were not viable past the first postnatal month (n=8). Nonetheless, consistent with our previous findings, P14 *caMek1*, *Nkx2.1:Cre, Ai9* mice displayed a reduction in PV^+^/RFP^+^-CIN density (Figure S2E-L, M). In summary, our data indicate that the establishment of PV-CIN number is cell-autonomously vulnerable to enhanced MEK1 signaling, while SST-CIN number is not altered.

### Presumptive mutant PV-CINs in the embryonic subpallium undergo apoptosis

Our analyses of RFP^+^ CINs in P3.5 *caMek1, Slc32A1:Cre* mice yielded a significant reduction in CIN density, thus, we hypothesized that gain-of-function MEK1 signaling disrupted embryonic processes necessary to establish CIN number. Indeed, examination of RFP^+^ CIN density in the E17.5 cortical plate also revealed fewer CINs in *caMek1, Slc32A1:Cre* embryos (Figure S3A-D). We examined markers of neuronal death during mid-neurogenesis in the *caMek1, Slc32A1:Cre, Ai9* subpallium. Immunolabeling for the apoptotic marker cleaved caspase 3 (CC3) revealed colocalization of CC3 with some RFP^+^ neurons within the mantle zone of the ganglionic eminences in E13.5 mutant, but not control, embryos (Figure 3A-E, F). CC3^+^/RFP^+^ neurons also presented with condensed, pyknotic nuclei (Figure 3G-N). We also observed CC3^+^/RFP^+^ cells with pyknotic nuclei in *caMek1*, *Nkx2.1:Cre, Ai9* mantle zone (Figure 3O-Q). No apoptotic cells were observed in the mutant ganglionic eminence VZ or the cortical migratory streams. Analysis of recombined GABAergic neuron density in the dorsal striatum did not reveal a significant difference from controls, suggesting that the loss of PV-CINs is not due to altered CIN migratory trajectory. Together, our results suggest reduced PV-CIN density in the postnatal cortex is due to the death of a subset of migrating CINs in the ganglionic eminence.

**Figure 3.**
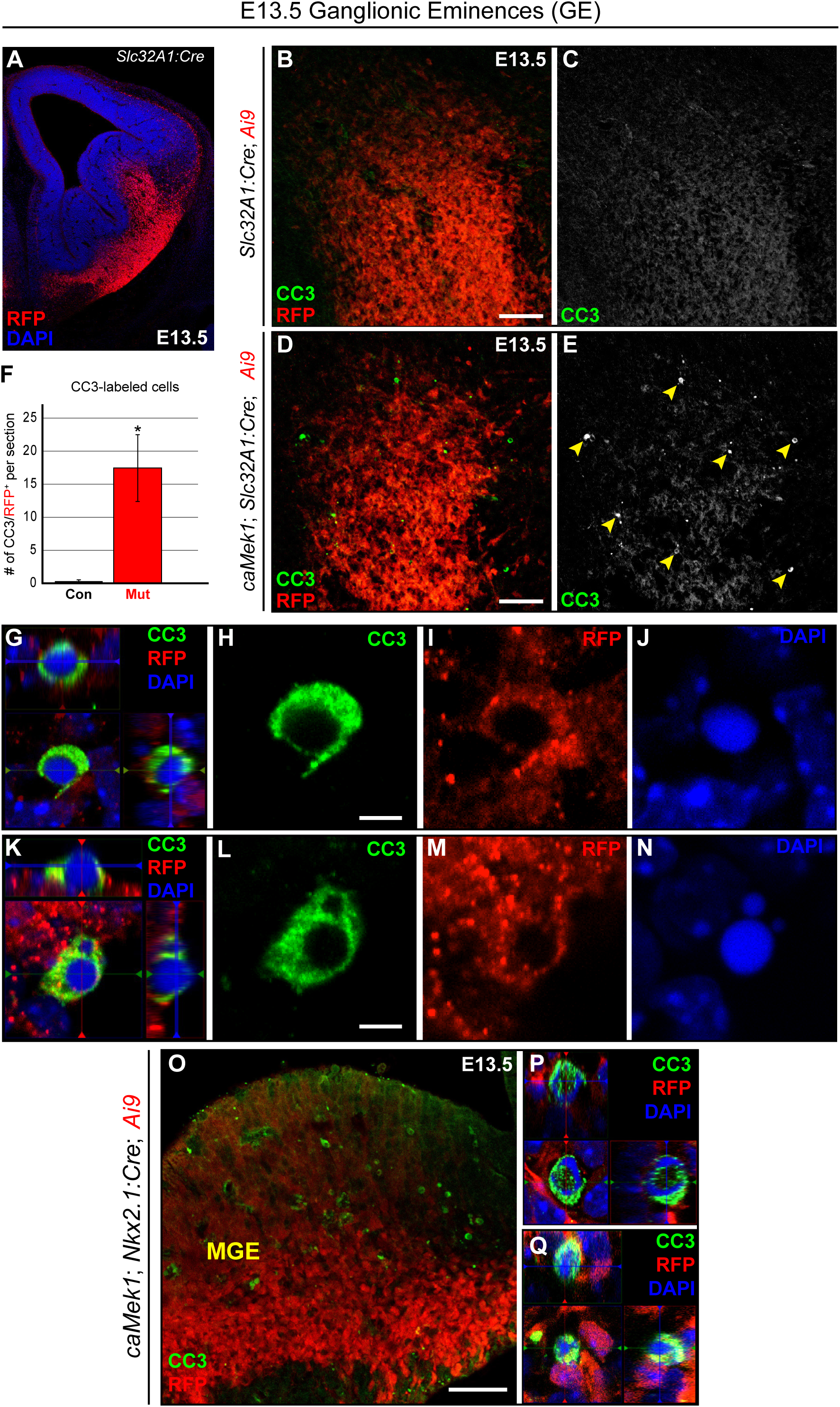
A subset of immature GABAergic neurons undergo cell death during mid-embryogenesis. **(A)** E13.5 coronal section of RFP-labeled CINs in the mantle zones of the *Slc32A1:Cre* subpallium during mid-embryogenesis. **(B-F)** Immunolabeling for cleaved caspase 3 (CC3) showed a significant increase in the number of apoptotic profiles in *caMek1, Slc32A1:Cre* mutants (D-E) as compared to controls (B-C) (quantification in **F**; n=3, mean ± SEM, * = p < 0.05). (Scale bar = 100 µm) **(G-Q)** Representative confocal z-stacks of CC3 labeled cells from *caMek1, Slc32A1:Cre* embryos (G-N, Scale bar = 2 µm) and *caMek1*, *Nkx2.1:Cre* embryos (O-Q) show clear colocalization with RFP and a condensed, pyknotic nuclear morphology. See also Figure S4.

### GABAergic-specific caMek1 promotes cortical hyperexcitability but does not significantly alter fast-spiking CIN electrophysiological properties

Nearly 40% of RASopathy individuals with mutations downstream of *RAS* experience seizures and epilepsy (Digilio et al., 2011; Rauen et al., 2013; Yoon et al., 2007). Whether MEK1 hyperactivation in GABAergic circuits mediates seizure activity in the RASopathies is unclear. We did not detect any signs of overt generalized tonic-clonic seizures in mutant mice while housed in home cages. We conducted a series of behavioral tests by first using the open field, then the elevated plus maze, and finally the social approach assay. No difference in global locomotor activity, anxiety-like behavior, or sociability could be detected with these tests (Figure S4). However, during the initial 60 sec of open field testing with 13 adult *caMek1, Slc32A1:Cre* mutants, two mutant mice exhibited increased head twitching, aberrant locomotor activity, and increased rearing (Supp. Video 1) and three mutant mice displayed periods of overt sudden behavioral arrest and motionless staring (Supp Video 2). These behaviors were not observed in any of the control mice utilized in this study. Consistent with these subtle impairments, subsequent re-analysis of the first 10 sec of the open field task revealed a significant reduction in distance traveled in *caMek1, Slc32A1:Cre* mutants, which was also observed in *caMek1, Dlx5/6:Cre* mutants.

We performed intracortical EEG recordings to directly assess cortical activity, which revealed spontaneous epileptiform-like discharges in three of six *caMek1, Slc32A1:Cre* adult mice, but not control mice (Figure 4A). These six mutants also exhibited a significantly reduced average threshold to seizure induction in response to pentylenetetrazol (PTZ) administration when compared to controls (Figure 4B). Seizures have been shown to increase the local expression of glial fibrillary acidic protein (GFAP) in astrocytes (Steward et al., 1992; Stringer, 1996). 3 of 3 newly generated and untreated *caMek1 Slc32A1:Cre* mice immunolabeled for GFAP exhibited clusters of GFAP-expressing astrocytes in the cortex consistent with local reactive astrogliosis near hyperexcitable regions (Figure 4C-F). Overall, some of the seizure-related phenotypes we observed were not completely penetrant, thus, these data indicate that MEK1 hyperactivation in CINs may be a potential risk factor for epilepsy in the RASopathies.

**Figure 4.**
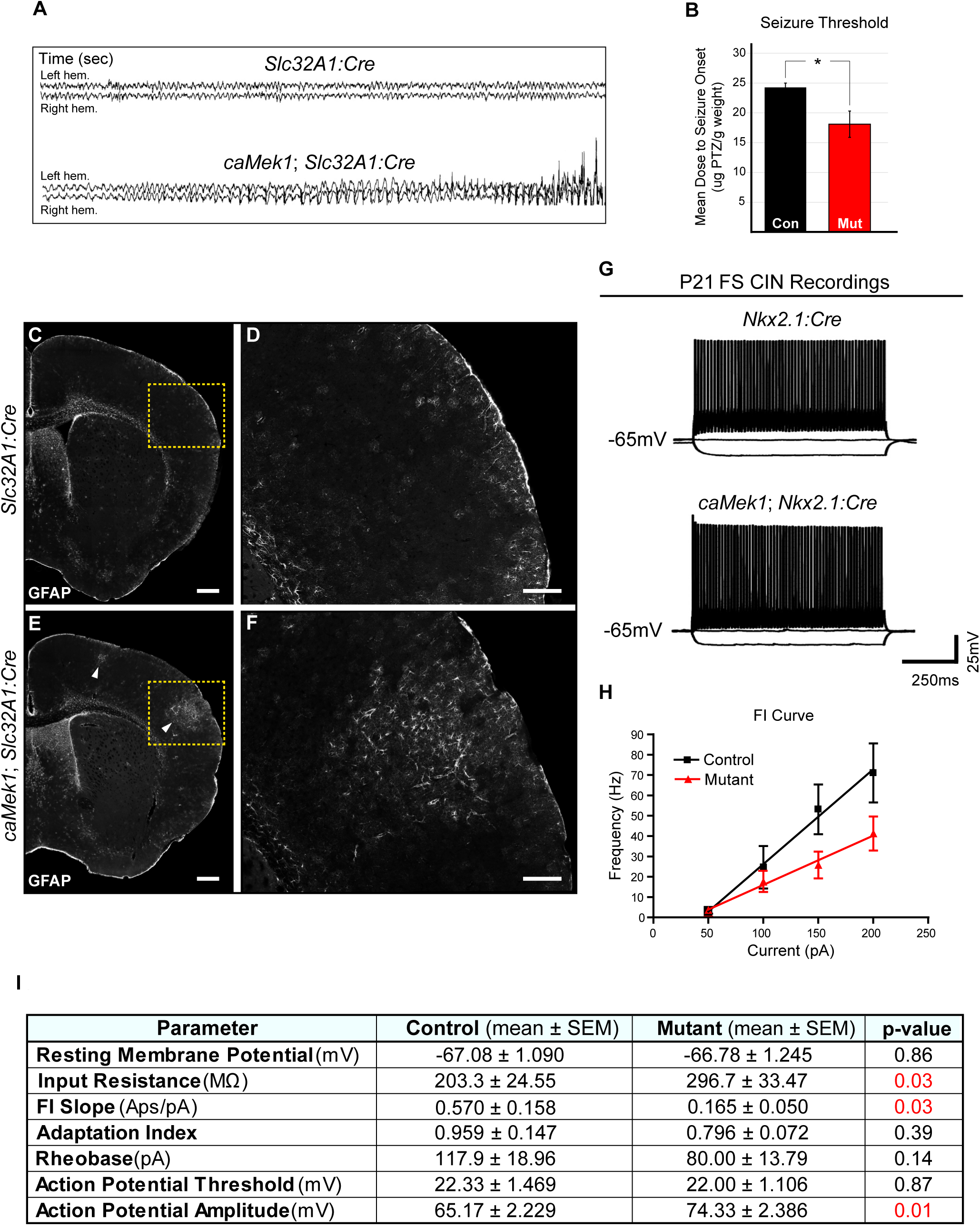
caMek1 Slc32A1:Cre CINs maintain typical fast-spiking properties, but a subset of mice exhibit seizure-like phenotypes. **(A)** Representative traces from forebrain-penetrating EEG revealed epochs of synchronous firing in 3 of 6 *caMek1 Slc32A1:Cre*, but not control mice. **(B)** Tail vein PTZ injections revealed a significant reduction in mean dose to seizure onset of PTZ (n = 6, mean ± SEM, * = p < 0.001). **(C-F)** caMek1 Slc32A1:Cre cortices display aberrant clusters of GFAP-labeled astrocytes (**E,** arrowheads, insets in **F**) that were not observed in controls (C-D) (n=3). (Scale bar = 100 µm) **(G)** Representative current clamp recordings of FS CINs in P21 *Nkx2.1:Cre Ai9* and *caMek1 Nkx2.1:Cre Ai9* mutant cortices. **(H)** Mutant CINs had a significantly reduced FI slope in comparison to controls (mean ± SEM, p < 0.05). **(K)** Summary table of FS CIN intrinsic properties. See also Figure S5.

PV-CINs provide a powerful source of inhibition in the cortex, firing action potentials at frequencies greater than 200Hz (Okaty et al., 2009). Fast-spiking (FS) physiology is due in part to the unique expression of the fast-inactivating potassium channel Kv3.1, which begins in the second postnatal week (Goldberg et al., 2011; Rosato-Siri et al., 2015; Rudy and McBain, 2001). To determine if hyperactive MEK1 signaling was sufficient to alter basic physiological properties of FS CINs, we performed whole-cell patch clamp recordings on *caMek1*, *Nkx2.1:Cre* mice at the end of the third postnatal week. Current clamp recordings of fluorescently-labeled CINs revealed that both control and mutant neurons retained their distinctive electrophysiological fast-spiking phenotype (Figure 4G, I) (Agmon & Connors, 2018; Anderson et al., 2010; McCormick et al., 1985). No significant differences were observed in resting membrane potential, adaptation index, rheobase, or action potential threshold, but a small increase in action potential amplitude was observed (Figure 4I). We did also detect a significant increase in FS CIN input resistance and a reduction in FI slope in *caMek1*, *Nkx2.1:Cre* mutant compared with *Nkx2.1:Cre* controls suggesting that mutants may have a reduction in the responsiveness and/or firing output of inhibitory neurons (Figure 5H-I). Overall, these data indicate that canonical electrophysiological features of fast-spiking CIN development were not altered by MEK1 hyperactivation. However, certain intrinsic properties exhibit subtle differences in mutant mice that might contribute to circuit-wide hyperexcitability.

**Figure 5.**
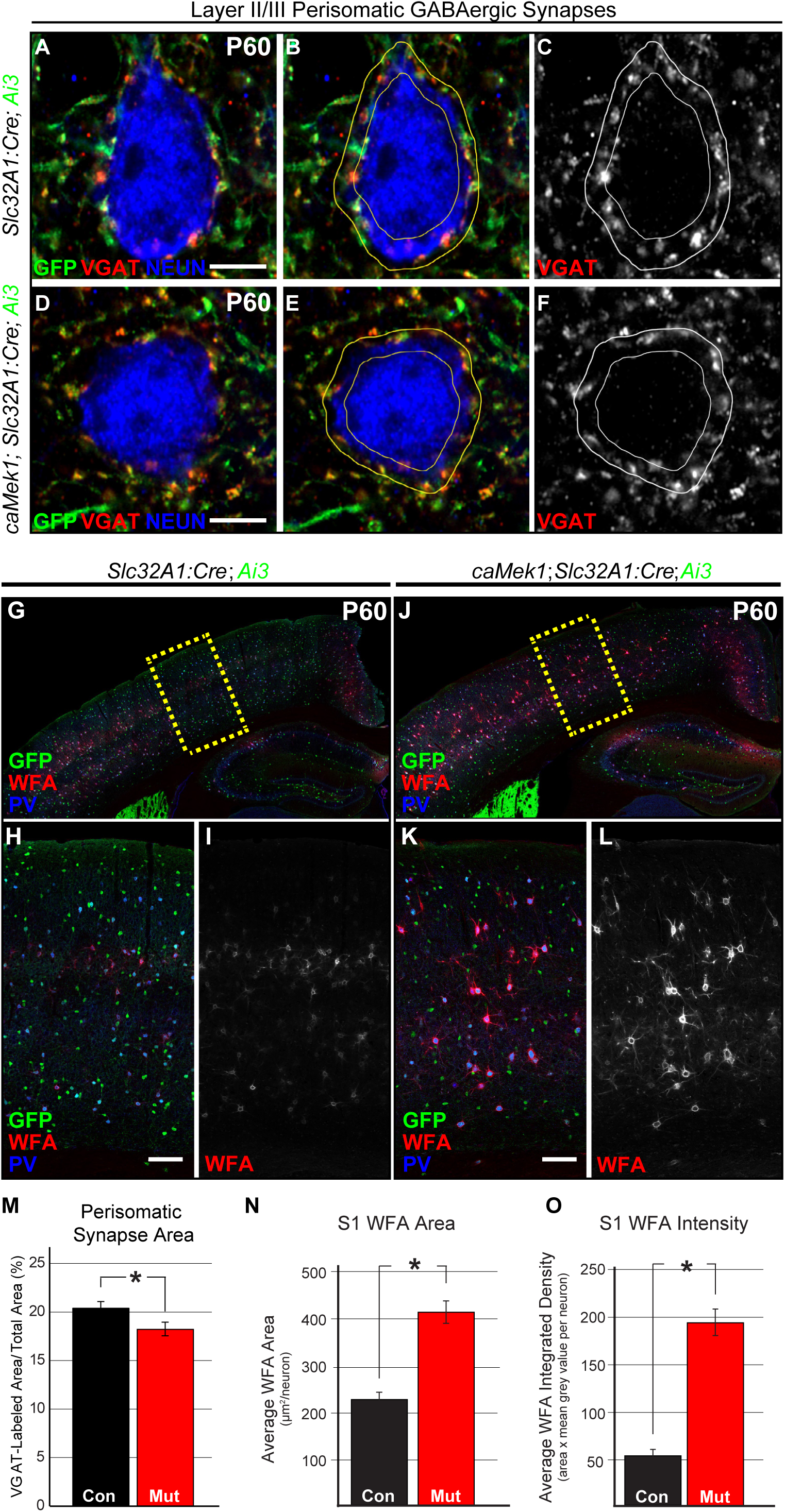
Reduced perisomatic synapse labeling in mutant cortices coincide with a substantial increase in PNN formation on PV-CINs. **(A-F)** Representative high-resolution confocal Airyscan images of triple immunolabeled cortical sections for Ai3/EYFP, VGAT, and NEUN. Excitatory neuron perisomatic domains were outlined and quantification of VGAT-labeled pixels revealed that mutants (D-F) have a significant reduction in the amount of perisomatic VGAT-labeling (Scale bar = 3 µm) in comparison to controls (A-C) (quantification in **M**; n = 48 control, 53 mutant neurons; mean ± SEM, * = p < 0.05). **(G-L)** P60 representative coronal sections of *Slc32A1:Cre Ai3* (G-I) and *caMek1 Slc32A1:Cre Ai3* (J-L) cortices immunolabeled for GFP, WFA, and PV. The WFA channel was imaged using the same acquisition settings for all samples. A significant increase in WFA-labeled area per neuron was detected in mutant cortices when compared to controls (quantification in **N**; n = 63 control, 54 mutant neurons, mean ± SEM, * = p < 0.001). Analysis of WFA-labeling intensity yielded a significant increase in integrated density in mutant CINs (quantification in **O**; n = 63 control, 54 mutant neurons, mean ± SEM, * = p < 0.001). (Scale bar = 100 µm).

### Inhibitory synapse formation and perineuronal net accumulation are altered by MEK1 hyperactivity

PV-expressing CINs preferentially innervate pyramidal cells, often forming synapses on the perisomatic domain (Chattopadhyaya et al., 2004; Chattopadhyaya et al., 2007). We assessed whether perisomatic VGAT-labeled synapses surrounding layer 2/3 PNs were diminished in adult *caMek1*, *Slc32A1:Cre, Ai3* mice. We found the extent of VGAT-immunolabeling in the perisomatic space of NEUN^+^/GFP^-^ PN soma was significantly reduced in mutant cortices when compared to controls (Fig. 5A-F, M). Interestingly, the area of VGAT-labeling in the surrounding neuropil, typically innervated by SST-CINs, was unchanged (Fig. S5A-F). These data show that PV-CIN inhibitory output is selectively vulnerable to caMEK1 signaling while SST-CINs are less affected.

PV-CINs selectively accumulate an extracellular structure called the perineuronal net (PNN) derived primarily from glial chondroitin sulfate proteoglycans (CSPGs). PNNs are essential to cortical development, restricting plasticity during the closure of critical periods and protecting PV-CINs from oxidative stress associated with a high frequency firing rate (Cabungcal et al., 2013; Hensch, 2005b). Reductions in PNN formation have been noted in multiple models of neurodevelopmental disorders that exhibit loss of PV-CINs (Bitanihirwe and Woo, 2014; Cabungcal et al., 2013; Krencik et al., 2015; Krishnan et al., 2015; Steullet et al., 2017). We utilized WFA-labeling to test whether PNN formation was reduced in adult *caMek1*, *Slc32A1:Cre, Ai3* mice (Figure 5H-M). Surprisingly, we found that surviving PV-CINs were WFA^+^ in mutants (Figure 5G-L). PNNs were also not detected on *Ai3*-expressing neurons that lacked PV-expression. Consistent with the larger somal size of mutant PV-CINs, the cross-sectional area of WFA-labeled profiles was also significantly increased (Figure 5G-L, N). Analysis of the quantitative level of WFA-labeling in mutant PV-CINs revealed a robust increase in PNN accumulation as compared to controls (Figure 5G-L, O). Mutant WFA-labeled CINs exhibited normal expression of 8-oxo-2’-deoxyguanosine (8-oxo-dg), a marker of DNA oxidation often altered in neurons with reduced PNNs (Figure S5 F-I) (Steullet et al., 2017). Collectively, MEK1 hyperactivation clearly increases PNN accumulation, but does not trigger ectopic PNN formation on GABAergic neurons lacking PV.

### caMek1, Slc32A1:Cre mice display delayed acquisition of FMI performance

Attention deficit hyperactivity disorder (ADHD) is associated with a significant proportion of RASopathy cases (Adviento et al., 2014; Garg et al., 2013; Green et al., 2017; Pierpont et al., 2018; Walsh et al., 2013). Altered prefrontal cortex (PFC) function has been implicated in ADHD (Gabay et al., 2018), and has been shown to contribute to cognitive deficits in a mouse model of Fragile X Syndrome (Krueger et al., 2011). Few studies have examined GABAergic contributions to PFC function in the context of RASopathies. As in sensory cortex, we noted a reduction in *Ai9*-expressing CINs in the PFC of mutant mice (Figure 6A-B, D-E). Interestingly, we found that PNs in the PFC exhibit reduced P-ERK1/2 expression in *caMek1, Slc32A1:Cre* mice (Figure 6A-F). These data indicate that MEK1 hyperactivation in developing CINs is sufficient to drive molecular abnormalities within specific cortical regions important for cognition.

**Figure 6.**
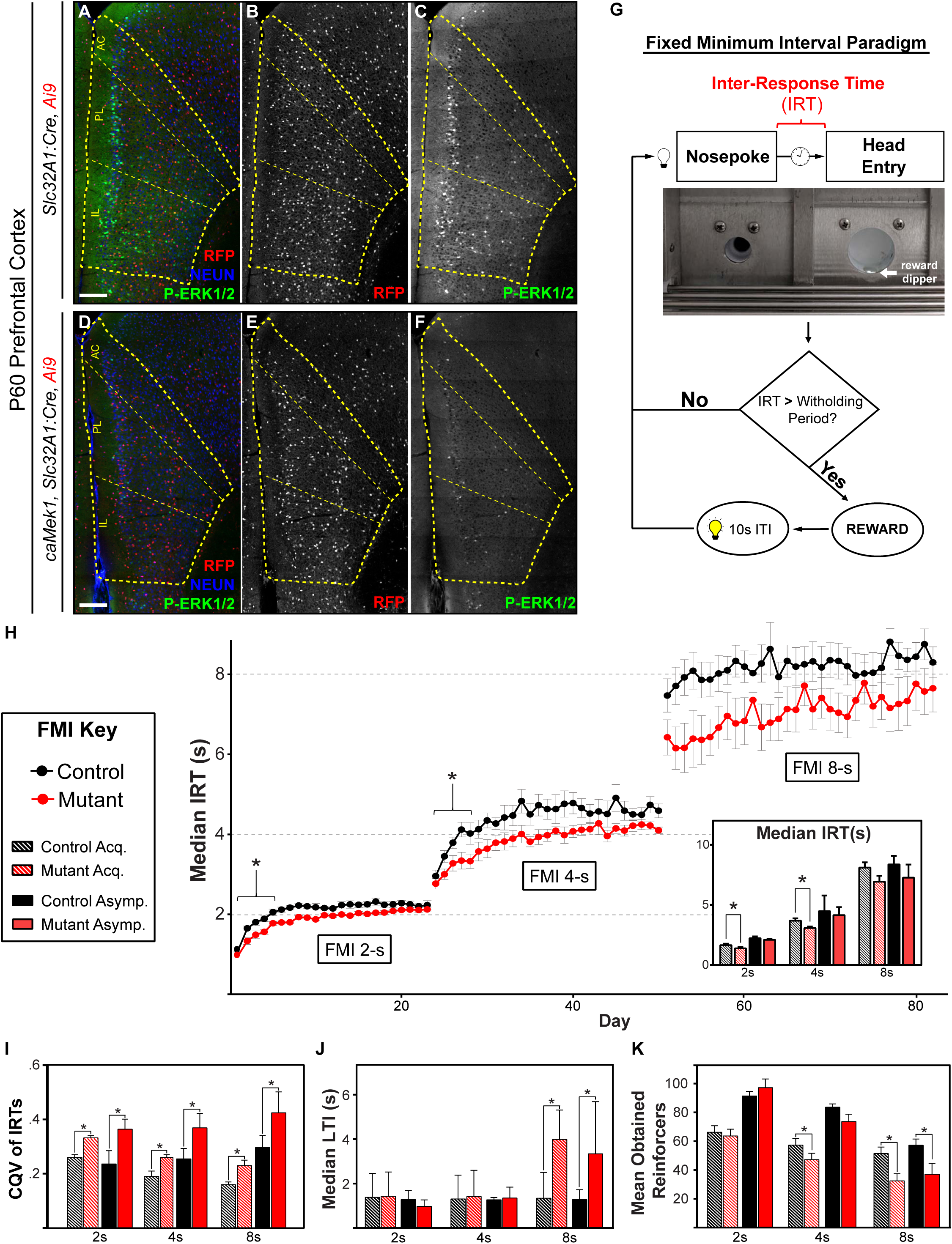
caMEK1 Slc32A1:Cre mice exhibit reduced behavioral response inhibition capacity. (**A-F)**. P-ERK1/2 labeling in PFC PNs is significantly reduced in mutants as compared to controls. Note the decrease in RFP^+^ CINs in the mutant PFC (E) relative to control (B) (n=3). (Scale bar = 100 µm) **(G)** Schematic of the Fixed-minimum Interval (FMI) task. **(H)** Mutant mice had a significant reduction in mean median IRT during FMI acquisition in 2s and 4s schedules (n=12, mean ± SEM, * = p < 0.05). **(I)** Mutant mean CQV of IRTs during both acquisition and asymptotic phases was significantly increased in 2s, 4s, and 8s FMI schedules (mean ± SEM, * = p < 0.05). **(J)** Median acquisition and asymptotic LTI was significantly increased in the FMI 8s, but not in the 2s and 4s schedules (mean ± SEM, * = p < 0.05). **(K)** Mutant mice had a reduction in mean acquisition ORs at 4s and a significant reduction in both mean acquisition and asymptotic ORs during the 8s FMI (mean ± SEM, * = p < 0.05).

Individuals with ADHD often exhibit structural changes in the PFC, which appear to be involved in the inhibition of reinforced responses (Seidman et al., 2006). To examine response inhibition directly, we utilized a fixed minimum interval (FMI) test, a timing-based task that requires animals to withhold a response for a fixed period. This paradigm is perhaps more favorable than the five-choice serial reaction time task (5-CSRTT) and differential reinforcement of low rates task (DRL), because its self-paced design dissociates response inhibition capacity from motivational aspects of behavior (Bizarro et al., 2003; Doughty and Richards, 2002; Hill et al., 2012; Watterson et al., 2015). Here, adult control and *caMek1, Slc32A1:Cre* mice were trained to initiate trials via a nose-poke which resulted in the presentation of sweetened condensed milk in the reward receptacle. Mice were then placed on an FMI schedule, where a time delay between the initiating nose-poke and the availability of reinforcement in the reward receptacle was implemented (Figure 8G). Reward was delivered only if the time between the initiating nose-poke and attempt to obtain reward (inter-response time, or IRT) exceeded a pre-determined withholding period. If mice prematurely accessed the reward receptacle, no reward was delivered.

Following initial training on a FMI with a very short (0.5s) response-withholding period, we measured mouse performance when the withholding period was extended to 2s, 4s, and finally, 8s. We observed a main effect of FMI schedule irrespective of genotype, such that IRTs increased as the FMI withholding period increased (F (2, 62) = 535.12, p < 0.01). Importantly, mutants showed clear evidence of impaired acquisition of the FMI task. We found a main effect of genotype on the mean median IRT during the first 5 days of each FMI schedule (acquisition period), in which mutant mice had relatively lower IRTs compared to control mice (F (1, 62) = 18.73, p < 0.01) (Figure 6H). In further support of reduced response inhibition capacity, mutant mice exhibited increased variability in their IRTs as measured by the coefficient of quartile variation (CQV) during acquisition (F(1, 62) = 31.73, p < 0.01) and asymptotic performance (defined as the last five days of the FMI) (F(2, 62) = 5.055, p < 0.001) across all schedules (Figure 6I). Median IRTs during the asymptotic phase in mutants and controls were not statistically different in any schedule (Figure 6H inset). Thus, these data suggest that mutant mice are capable of learning to inhibit reinforced responses for up to 4 s but show a significant delay in acquiring this capability.

Due to *Slc32A1:Cre*-mediated recombination within subcortical circuitry, it is possible that altered reward pathway activity influenced FMI performance. The latency to initiate (LTI) a trial provides a measure of motivation; for example, rats administered amphetamine show a reduction in LTI in a related task (Rojas-Leguizamón et al., 2018). However, we noted that mutants did not differ from controls in the median LTI at 2s and 4s, indicating that motivation to obtain rewards was not significantly altered between conditions (Tukey’s b post-hoc test −2s: t(21) = 1.39, p=0.18; 4s: t(21) = −0.29, p=0.77) (Figure 6J). We found that during the 8s FMI, mutant mice exhibited a statistically significant increase in LTI (t(20) = 2.43, p < 0.05). This apparent loss of motivation is likely due to the fact that the mean median asymptotic IRT did not reach the 8s criterion even after 32 days of testing (control: 8.45s ± 1.04; mutant: 7.26s ± 1.09) and is also consistent with the statistically significant reduction in mean obtained reinforcers at the 8s FMI (Figure 6K). Collectively, our data indicate that altered GABAergic circuitry regulates acquisition of response inhibition capacity in mice and may contribute to ADHD phenotypes associated with the RASopathies.

## Discussion

Here, we show that GABAergic neuron-autonomous MEK1 hyperactivation causes the death of a subset of immature GABAergic neurons in the embryonic subpallium and is associated with a selective reduction in PV-CIN density, but not SST-CINs, in adulthood. We observed a significant reduction in perisomatic GABAergic synapses on layer 2/3 PNs and forebrain hyperexcitability, but a surprising increase in the extent of PNN accumulation in mutant PV-CINs. While mutants displayed relatively normal performance in assays of locomotion, sociability, and anxiety, we found notable defects in acquisition of behavioral response inhibition capacity, which has been linked to ADHD. These data suggest that GABAergic neuron-autonomous MEK1 hyperactivation selectively regulates embryonic PV-CIN survival and is an important contributor to seizure risk and cognitive deficits in the RASopathies.

While expression of ERK/MAPK pathway components is widespread, our findings reinforce the notion that expression levels are variable, and activation of this cascade is highly cell-type dependent. Cell-specific transcriptomic experiments have reported that RNA levels of *Mapk1*/*Erk2* and *Map2k1/Mek1*, but not *Mapk3/Erk1* or *Map2k2/Mek2*, are lower in CINs relative to PNs (Mardinly et al., 2016). We have extended these findings to show that protein levels of MAPK1/ERK2 and MAP2K1/MEK1 are typically lower in CINs than in surrounding PNs. Reduced expression of pan-ERK/MAPK components may contribute to the relatively low levels of phosphorylated ERK1/2 in CINs. Past work has shown that the experience-dependent transcriptional response in V1 PV-CINs is significantly smaller relative to PNs (Hrvatin et al., 2018). A more stringent degree of ERK/MAPK recruitment in CINs might contribute to reduced activity-dependent transcription. Further analysis of activity-dependent responses in ERK/MAPK-mutant mice may assist in defining how CIN-specific functional properties are encoded (Tyssowski et al., 2018).

Defects in GABAergic circuitry have been implicated in the pathogenesis of Rett, Fragile X, schizophrenia and many other neurodevelopmental diseases (Chao et al., 2010; Cui et al., 2008; Steullet et al., 2017). Reduced PV-CIN number is often observed, however, the mechanism of loss is poorly understood. We show that MEK1 hyperactivation drives the GABAergic-neuron autonomous activation of caspase-3 and death of a subset of immature neurons in the embryonic ganglionic eminences. The selective reduction in PV-CIN density in the postnatal cortex suggests these early dying neurons were committed to the PV lineage. The death of this specific subset of GABAergic neurons occurs much earlier than the typical period of programmed cell death for CINs (Denaxa et al., 2018; Southwell et al., 2012). Notably, *caMek1* expression in cortical excitatory neurons is not associated with significant neuronal loss during development (Nateri et al., 2007; Xing et al., 2016). Though ERK/MAPK typically acts as a promoter of cell survival, apoptotic death by sustained ERK/MAPK activity has been described in certain contexts (Cagnol and Chambard, 2010; Martin and Pognonec, 2010). It will be important to evaluate whether treatment with pharmacological MEK1/2 inhibitors or antioxidants during embryogenesis is capable of sustained restoration of CIN number in *caMek1*, *Slc32A1:Cre* mice. PV-CIN sensitivity to MEK1 hyperactivation may not only be an important factor in RASopathy neuropathology, but could be a relevant mechanism in other conditions that involve indirect activation of ERK/MAPK signaling during embryogenesis, such as schizophrenia, Fragile X Syndrome, or prenatal stress (Fowke et al., 2018).

Recent scRNAseq analyses suggest the mature transcriptional signature of cardinal CIN subtypes is not fully specified until CINs migrate into the cortex (Mayer et al., 2018; Paul et al., 2017; Sandberg et al., 2018). MEF2C, a known substrate of ERK1/2 and p38 signaling, was identified as a transcription factor expressed early in the presumptive PV lineage, the deletion of which also leads to the selective reduction of PV-CINs (Mayer et al., 2018). Thus, regulation of MEF2C could mediate early effects of caMEK1 signaling on GABAergic neuron development. While further research is necessary, the selective vulnerability of presumptive PV-CINs to hyperactive MEK1 signaling may not be dependent upon a specific downstream transcriptional target. Compared to transcriptional networks, less is known regarding the function of many post-translational modifications during CIN specification. Our data hint at selective roles for kinase signaling networks at an early stage of CIN lineage differentiation.

Despite the effect of caMEK1 on early GABAergic neuron survival, the physiological maturation of mature *caMek1*-expressing PV-CINs was not significantly impeded. Surviving PV-CINs retained a characteristic fast-spiking signature with only minor differences in intrinsic electrophysiological parameters. We noted a modest, but statistically significant decrease in perisomatic inhibitory synapse number on cortical PNs in mutant mice. As might be expected, a subset of mutant animals exhibited forebrain hyperexcitability and sudden behavioral arrest similar to that reported in animal models of mild seizures. Overexpression of a similar *caMek1* mutation with *CamKII:Cre* has also been shown to cause seizure-like activity (Nateri et al., 2007). Overall, our findings indicate that MEK1 hyperactivation in GABAergic neurons could increase the risk of epilepsy seen in RASopathies.

The PNN is a critically important structure involved in the maturation of cortical circuitry with an important role in protecting PV-CINs from oxidative stress and limiting synaptic plasticity (Cabungcal et al., 2013; Hensch, 2005a; Hensch, 2005b; Morishita et al., 2015). Mouse models of schizophrenia, Fragile X, and ASDs often exhibit reduced PV-CIN number and typically display a reduction in PNN formation (Steullet et al., 2017). In contrast to these disorders, PNNs appear to respond differently to RASopathy mutations. PV-CINs accumulate extracellular PNNs derived primarily from astrocyte-produced CSPGs (Galtrey and Fawcett, 2007; Sorg et al., 2016). RASopathic astrocytes upregulate secreted ECM-associated CSPGs and promote an increase in the extent of PNN accumulation around PV-CINs (Krencik et al., 2015). Our data is the first to indicate a PV-CIN autonomous role for enhanced PNN accumulation in response to MEK1 hyperactivation. It is thought that increased PNN accumulation on PV-CINs limits the plasticity of cortical regions (Pizzorusso et al., 2002). Modification of PNN levels may serve as a useful therapeutic strategy for the impaired cognitive function and intellectual disability frequently reported in RASopathy individuals (Tidyman and Rauen, 2016).

In addition to intellectual disability, ADHD is frequently diagnosed in Noonan Syndrome and NF1, two common RASopathies (Johnson et al., 2019; Miguel et al., 2015; Pierpont et al., 2015). Abnormal PFC function has been linked to ADHD (Seidman et al., 2006). Interestingly, we detected significantly reduced P-ERK1/2 levels in PFC PNs of mutant mice, suggesting that RASopathic CINs may alter the global development and function of this brain region. GABAergic signaling is known to be necessary for cortical circuit maturation (Cancedda et al., 2007). We further examined ADHD-related behavioral phenotypes in *caMek1, Slc32A1:Cre* mice by assessing behavioral response inhibition capacity with a fixed-minimum interval (FMI) based task (Rojas-Leguizamón et al., 2018; Watterson et al., 2015). We detected significant deficits in the acquisition of response inhibition dependent behaviors in mutant mice relative to controls. It is plausible that FMI defects in caMEK1 mutants is due to the reduced plasticity of PFC GABAergic circuitry in response to heightened levels of PNN or GABAergic-dependent changes in PN development. These data show that GABAergic-directed MEK1 hyperactivation is sufficient to drive deficits in behavioral response inhibition possibly associated with ADHD.

Mutations in ‘upstream’ RASopathy genes modulate a much broader set of downstream cascades when compared to mutations in *Raf* or *Mek1/2*. *Nf1*, *Ptpn11/Shp2*, and *Syngap1* mutations result in a complex constellation of cellular changes, some of which depend upon ERK/MAPK modulation, whereas others involve different signaling cascades (Anastasaki and Gutmann, 2014; Brown et al., 2012). In combination with the findings of (Angara et al., 2019, co-submitted), it is clear that PV-CIN development is particularly sensitive to convergent signaling via NF1 and ERK/MAPK. Additional studies of human samples will be necessary to determine whether defective GABAergic circuits are a component of RASopathy pathogenesis. Collectively, our research suggests that hyperactivation of MEK1 in GABAergic neurons represents an important candidate mechanism for epilepsy and cognitive defects in RASopathic individuals.

## Acknowledgements

We would like to thank Johan Martinez, Sam Lusk, Julia Pringle, Anna Kreuger, Sarah Sparks, Katie Riordan, Danielle Gonzalez, Elise Bouchal, and Kimberly Holter for their technical contributions to this work. This research is supported by National Institute of Health grants R00NS076661 and R01NS097537 awarded to JMN, R01NS087031 awarded to TRA, and R01NS031768 to WDS.

## Author Contributions

Conceptualization, J.M.N, W.D.S, M.C.H, C.W.D, F.S., T.R.A.; Methodology, J.M.N, W.D.S, M.C.H, C.W.D, F.S., T.R.A., S.M., D.M.T.; Investigation, J.M.N, W.D.S, M.C.H, L.T.H., K.N., G.R.B., S.S., N.R.F., K.P.R., T.A.G., G.L., M.F.O., C.W.D, F.S., T.R.A., S.M., D.M.T.; Writing – Original Draft, M.C.H., J.M.N., C.W.D., T.A.G., F.S., T.R.A., S.M., M.F.O., L.T.H., K.N.; Writing – Review & Editing, M.C.H., J.M.N., G.R.B., S.S., N.R.F., K.P.R., D.M.T.; Funding Acquisition, J.M.N and W.D.S,; Resources, J.M.N, T.R.A., W.D.S., D.M.T., M.F.O., F.S.; Supervision, J.M.N., W.D.S., D.M.T., T.R.A., F.S.

## Declaration of Interest

The authors declare no competing interests.

## Methods

### Mice

All transgenic mice were handled and housed in accordance with the guidelines of the Institutional Animal Care and Use Committee at Arizona State University, the University of Arizona, and Barrow Neurological Institute. Mice were kept on a daily 12-hour light-dark cycle and were fed *ad libitum*. *Slc32A1:Cre*^+/+^, *Dlx5/6:Cre^+/-^*, or *Nkx2.1:Cre*^+/-^ mice were crossed with *CAG-lox-STOP-lox-Mek1^S217/221E+/-^* (caMEK1) mice to generate mutants expressing *caMek1* in Cre-expressing cell types (Table S1). *CaMek1* mice were kindly provided by Dr. Maike Krenz and Dr. Jeffrey Robbins. Littermates expressing Cre-Recombinase were utilized as controls for most experiments, unless otherwise indicated. Cre-dependent tdTomato (*Ai9*) or eYFP (*Ai3*) strains were used to endogenously label Cre-expressing cells for visualization purposes. Genomic DNA was extracted from tail or toe samples for standard genotyping by PCR using the following primer combinations: (listed 5’-3’): Cre – TTCGCAAGAACCTGATGGAC and CATTGCTGTCACTTGGTCGT to amplify a 266 bp fragment; *caMek1^S217/221E^*-GTACCAGCTCGGCGGAGACCAA and TTGATCACAGCAATGCTAACTTTC amplify a 600 bp fragment; *Ai3/Ai9* – AAGGGAGCTGCAGTGGAGTA, CCGAAAATCTGTGGGAAGTC, ACATGGTCCTGCTGGAGTTC, and GGCATTAAAGCAGCGTATCC amplify a 297 bp wt Rosa26 segment and a 212 bp *Ai3*/*Ai9* allele.

### Tissue Preparation and Immunostaining

Mice of the appropriate postnatal age were anesthetized and transcardially perfused with PBS followed by cold 4% PFA in PBS. Brains were dissected, post-fixed at 4°C, and sectioned with a vibratome or cryopreserved with 30% sucrose and sectioned with a cryostat. Free-floating sections were incubated in primary antibody solution consisting of 1X PBS with 0.05 - 0.2% Triton and 5% Normal Donkey Serum (NDS). Sections were then incubated in species-specific, fluorescently-conjugated secondary antibodies in blocking solution overnight. For embryonic sections, timed-bred embryos were collected at the appropriate embryonic age, immersion fixed in cold 4% PFA in 1X PBS and cryopreserved in serial sucrose concentrations (15%, 25% in 1X PBS) until fully infiltrated. Embryonic brain sections were cryosectioned and directly mounted onto Fisher Superfrost Plus slides. Sections were gently rinsed in 1X PBS 0.05% Triton, incubated in blocking solution (1X PBST 0.05% Triton and 5% NDS) and incubated overnight in primary antibody prepared in blocking solution. The primary antibodies used in these experiments were: goat anti-parvalbumin (Swant, 1:1000), rabbit anti-somatostatin (Peninsula, 1:1000), biotin-conjugated WFA (Vector, 60ug/mL), chicken anti-GFP (Aves, 1:1000), chicken anti-RFP (Rockland, 1:1000), rabbit anti-P-ERK (Cell Signaling, 1:1000), rabbit anti-MEK1 (Abcam, 1:1000), rabbit anti-ERK2 (Abcam, 1:1000), rabbit anti P-ERK1/2 (Cell Signaling, 1:1000), mouse anti-NEUN (Millipore 1:1000), rabbit anti-cleaved caspase 3 (Cell Signaling, 1:1000), rabbit anti-GFAP (Abcam 1:1000), rabbit anti-VGAT (Synaptic Systems, 1:1000), and mouse anti-8-oxo-DG (R&D Systems, 1:1000) (Table S2). Tissue was then washed in 1X PBS 0.05% Triton and incubated in fluorescently conjugated secondary antibody solution before rinsing and cover-slipping for microscopic analysis. Alexa-Fluor 488, 568, and 647 conjugated anti-rabbit, anti-goat, and anti-chicken antibodies were diluted to 1:1000 in 1X PBS 0.05 – 0.2% Triton and 5% NDS. Streptavidin-conjugated fluorophores were used to visualize WFA labeling. Representative images were collected on a Zeiss (LSM710 & LSM800) laser scanning confocal microscope and optimized for brightness and contrast in Adobe Photoshop.

### Image Analysis

Images of at least three anatomically matched sections that include a brain region of interest were quantified for labeled cell density by observers blind to genotype. For estimating labeled cell density in the cortex, a column spanning all cortical layers was defined, the cross-sectional area measured, and the number of labeled cells was assessed. The proportion of cells co-labeled with Cre-dependent fluorescent reporters was also determined for select experiments. Quantification of cellular labeling was averaged across all images collected from an individual mouse. At least three mice were collected for each genotype and results were analyzed using Student’s t-tests unless indicated otherwise.

We quantified the extent of inhibitory synapse labeling in the perisomal domain of excitatory neurons from confocal images of VGAT/NEUN/GFP co-labeled sections. Confocal images were collected using optimal Airyscan settings for a 63x 1.4 NA objective on a Zeiss LSM800 with the same acquisition parameters, laser power, gain, and offset for VGAT detection. NEUN^+^/GFP^-^ neurons in S1 layer 2/3 with a pyramidal morphology and residing 5-10μm from the tissue section surface were randomly selected by a blinded observer. NEUN^+^ soma were outlined in Photoshop and a ring 1.8μm in thickness was then established to specify the perisomatic space. VGAT-immunolabeling from perisomatic regions of interest were imported into ImageJ where a moment-preserving autothreshold algorithm, “Moments”, was utilized to define the total area of perisomatic VGAT-labeling in an unbiased manner. The perisomal VGAT-labeled area was then normalized to the total perisomatic area for that neuron. A total of 48 control and 53 mutant neurons from three different mice were analyzed. A similar approach was utilized to quantify VGAT labeling in areas enriched in dendrites by analyzing randomly selected regions of the layer 2/3 neuropil that did not incorporate any NEUN-labeled soma.

### EEG Recordings and Seizure Threshold Assessment

Adult *caMek1*, *Slc32A1:Cre* mutant and *Slc32A1:Cre* control mice were assessed for epileptiform activity with bilateral 175μm tungsten wires implanted in the forebrain. After recovery from electrode implantation, mice were connected to suspended EEG leads, housed individually, and monitored daily in home cages for seizure-like activity using a 128 channel Natus Medical EEG machine. EEG recordings were examined for synchronous firing between hemispheres and representative epileptiform traces were acquired. Following intracranial recording, mice were injected with the seizure inducing compound, Pentylenetetrazol (PTZ; Sigma P6500). Mice were gently restrained and the tail vein was intravenously injected with 0.34ml/min of 5mg/mL PTZ in 0.9% saline 10USP heparin by automated pump. Initial onset of seizure was defined as the first sign of involuntary movement by an observer blinded to genotype. Time to seizure was recorded and PTZ μg/g of body weight was calculated.

### Slice electrophysiology

*caMek1^+/-^, Nkx2.1:Cre^+/-^, Ai9*^+/-^ mutant and *Nkx2.1:Cre^+/-^, Ai9*^+/-^control mice were sacrificed between postnatal day 21 to 24 and used for the in vitro slice electrophysiology. Brain slicing was performed as reported previously (Nichols et al., 2018). In brief, mice were deeply anesthetized by isoflurane inhalation before decapitation. Brains were quickly removed and the coronal slices (350 µM) of the somatosensory cortex were produced on a vibratome (VT 1200; Leica, Nussloch, Germany) in fully oxygenated (95% O_2_, 5% CO_2_), ice-cold artificial cerebral spinal fluid (aCSF) containing (in mM): 126 NaCl, 26 NaHCO_3_, 2.5 KCl, 10 glucose, 1.25 Na_2_H_2_PO_4_·H_2_O, 1 MgSO_4_·7H_2_O, 2 CaCl_2_·H_2_O, pH 7.4. The slices were incubated in the same aCSF at 32°C for 30min before being allowed to recover at room temperature for an additional 30 min before patch clamp recordings were started.

After recovery, slices were transferred into recording chamber and perfused continuously with aCSF of 32°C at a rate of 1-2 ml/min. Then whole-cell patch clamp recordings were performed on tdTomato-positive fast-spiking (FS) interneurons in the somatosensory cortex layer V/VI (L5/6) by using an Axon 700B amplifier. The FS neurons were identified by lack of an emerging apical dendrite and their intrinsic firing response to current injection (Agmon & Connors, 2018; Anderson et al., 2010; McCormick et al., 1985). Clampex 10.6 (Molecular Devices) was used to collect data and pipettes (2-5 MΩ) were pulled from borosilicate glass (BF150-110-10, Sutter Instruments) by using sutter puller (Model P-1000, Sutter Instruments), filled with an internal solution that contains (in mM): 135 K-Gluconate, 4 KCl, 2 NaCl, 10 HEPES, 4 EGTA, 4 Mg ATP and 0.3 Na Tris. The stability of the recordings was monitored during the experiments, and only the recordings with the series resistances (R_s_) less than both 25 MΩ and 20% of the membrane resistances were chosen for analysis. For the input resistance calculation, the steady plateau of the voltage responding to the current input of −50 pA step with 1 s duration was used and intrinsic parameters were measured as previously reported (Nichols et al., 2018). Adaptation index was calculated as the ratio of the 1^st^ interspike interval over the last (i.e. F_1st ISI_/F_last ISI_). The frequency (F) – current (I) slope was calculated as the number of induced action potentials (APs) divided by the current step (number of APs at 150pA-number of APs at 100pA)/(150pA-100pA). Unpaired Student’s t-test and two-way ANOVA with Bonferroni post hoc tests were used for statistical analysis.

### Behavioral Testing

#### Open Field Testing

The open field test was used to test voluntary locomotor capabilities and anxiety-like behavior. The apparatus consisted of a 40×40cm arena enclosed by 30cm high opaque walls. A single 60W bulb was positioned to brightly illuminate the center of the chamber with dim lighting near the walls. Mice were placed into the apparatus and recorded for a total of 10 minutes. Video data were analyzed for total distance traveled and time spent in the center quadrant.

#### Elevated Plus Maze

The elevated plus maze was constructed from black polycarbonate, elevated 81cm off the ground, and oriented in a plus formation with two 12×55cm open arms and two 12×55cm closed arms extending from an open 12×12cm center square. Closed arm walls were 40cm high extending from the base of the arm at the center square. The apparatus was lit with a 60W bulb with light concentrated on the center square. At the beginning of the trial, mice were placed in the center square, facing the south open arm, and recorded while freely exploring for 5 minutes.

#### Social Approach Assay

The social approach apparatus was made of transparent plexiglass and contained three 20×30×30cm chambers (total dimensions 60×30×30cm) connected by open doorways. Prior to experimental social trials, mice were habituated to the apparatus and allowed to freely explore all three chambers for 5 minutes. At the end of the 5 minutes, mice were removed and placed in their home cage. A sex- and age-matched stimulus mouse was then placed into a small empty cage in chamber 1 of the apparatus. The experimental mouse was reintroduced to the center chamber (chamber 2) of the apparatus and recorded while freely exploring for 10 minutes. The time spent in the chamber with the stimulus mouse (chamber 1) or the empty chamber (chamber 3) was then measured.

#### Fixed Minimum Interval (FMI)

Twenty-four adult mice (12 *Slc32A1:Cre* mice: 5 males, 7 females; 12 *caMek1*, *Slc32A1:Cre* mice: 6 males, 6 females) were kept on a 12-hour reverse light-dark cycle. Animals had free access to water in their home cages, but access to food was gradually reduced in the week prior to behavioral training, where 1 hr of food access was provided 30 min after the end of each daily training session. Body weights were maintained such that mice lost no more than 15% of starting body weight. Behavioral testing was conducted in eight MED Associates (St. Albans, VT, USA) modular test chambers (240 mm length × 200 mm width × 165 mm height; ENV-307W). Each chamber was enclosed in a sound- and light-attenuating cabinet (ENV-022V) equipped with a fan for ventilation that provided masking noise of approximately 60dB (ENV-025-F28). The back wall and hinged front door of each chamber were made of Plexiglas. The side walls of the chamber were made of aluminum, and the right wall contained the manipulanda and reward receptacle. The floor was composed of thin metal bars. A circular reward receptacle was positioned in the center of the front panel and equipped with a head entry detector (ENV-302HD), a liquid dipper (ENV-302W-S), and a yellow LED (ENV-321W). The reward receptacle was flanked by a nose-poke device including an LED-illuminator (ENV-314M). The chamber was fitted with a house light (ENV-315W) at ceiling level above the back wall (ENV-323AW) and a 4.5kHz tone generator (ENV-323HAM). Experimental programs were arranged via a MED PC interface connected to a PC controlled by MED-PC IV software. All behavioral sessions were 30 min long, including a 3-min warm-up period during which no stimuli were activated.

##### Reinforcement Training and Autoshaping

Mice were first trained to obtain 0.1 cc of diluted sweetened condensed milk from the liquid dipper (the reinforcer) in the reward receptacle. Following the 3-min warmup period, a reinforcer was made available, followed by consistent reinforcer delivery at variable, pseudo-randomly selected inter-trial intervals (ITIs) for the remainder of the session (mean = 45 s). No stimuli were activated during ITIs. When the dipper was activated and a reinforcer was available, a 2.9-kHz tone, the head-entry LED, and the house light were turned on. The reinforcer remained available until it was obtained by the mouse, which deactivated the 2.9-kHz tone, the LED, and house light. The dipper remains activated for 2.5s after the mouse obtains the reinforcer. Following 5 sessions of reinforcement training, the procedure was modified for 5 autoshaping sessions which, in the last 8s of each ITI, the LED inside of the nose-poke device was turned on. The nose-poke LED was then turned off and reinforcement was delivered as described. If the mouse nose-poked the device during the time when the LED was on, it was turned off and reinforcement was delivered immediately. The autoshaping procedure was then modified for another 5 sessions such that reinforcement delivery was contingent upon a single nose-poke to the nose-poke device when its LED was illuminated and the ITI was reduced to 10s.

##### Fixed-Minimum Interval Training

Mice were then trained on the fixed-minimum interval (FMI) schedule. After the 3-min warmup period, the houselight was deactivated. A nose-poke (*initiating response*) activated the nose-poke LED and marked the beginning of the inter-response time (IRT). A subsequent head entry into the reward receptacle (*terminating response*) terminated the IRT. Reinforcement was delivered only if the IRT was longer than the criterion time, which was dependent upon the FMI schedule. IRTs shorter than the criterion time terminated without reinforcement, deactivated the nose-poke LED, and another trial could be immediately initiated. IRTs greater than or equal to the criterion time resulted in delivery of reward, deactivation of the nose-poke LED, a 2.5s duration 2.9kHz tone, and subsequent removal of the liquid dipper. Houselights were then activated for a 10s ITI, after which houselight deactivation indicated a new trial could be initiated via nose-poke. The time between the end of the ITI and the nose-poke initiating response was measured and termed the *latency to initiate* (LTI). All mice were initially trained on an FMI schedule with a criterion time of 0.5s (FMI 0.5s) until stability was achieved. The FMI 0.5s condition was implemented to acclimate mice to the task and is not used to evaluate response inhibition capacity. Performance was considered stable when a non-significant linear regression for mean median IRTs across 5 consecutive sessions was achieved, using a significance criterion of .05. Following stability on the FMI 0.5s schedule, subjects experience FMI 2s, 4s, and 8s. Each subject was trained to stability.

##### Data Analysis

Four parameters were tracked on a session-by-session basis: median latency-to-initiate trials (LTI), median inter-response time (IRT), the coefficient of quartile variation (CQV) of IRTs (difference between 1^st^ and 3^rd^ quartile divided by their sum), and the number of obtained reinforcers (ORs). The acquisition phase of each parameter was defined as the mean performance during the first five sessions of each schedule, while the asymptote was defined as the mean during the last five sessions. ANOVAs were conducted to assess statistical significance of time and genotype on FMI schedule and Student’s *t*-tests were conducted to examine parameter differences based on genotype.

## Supplemental Information

**Table S1.**
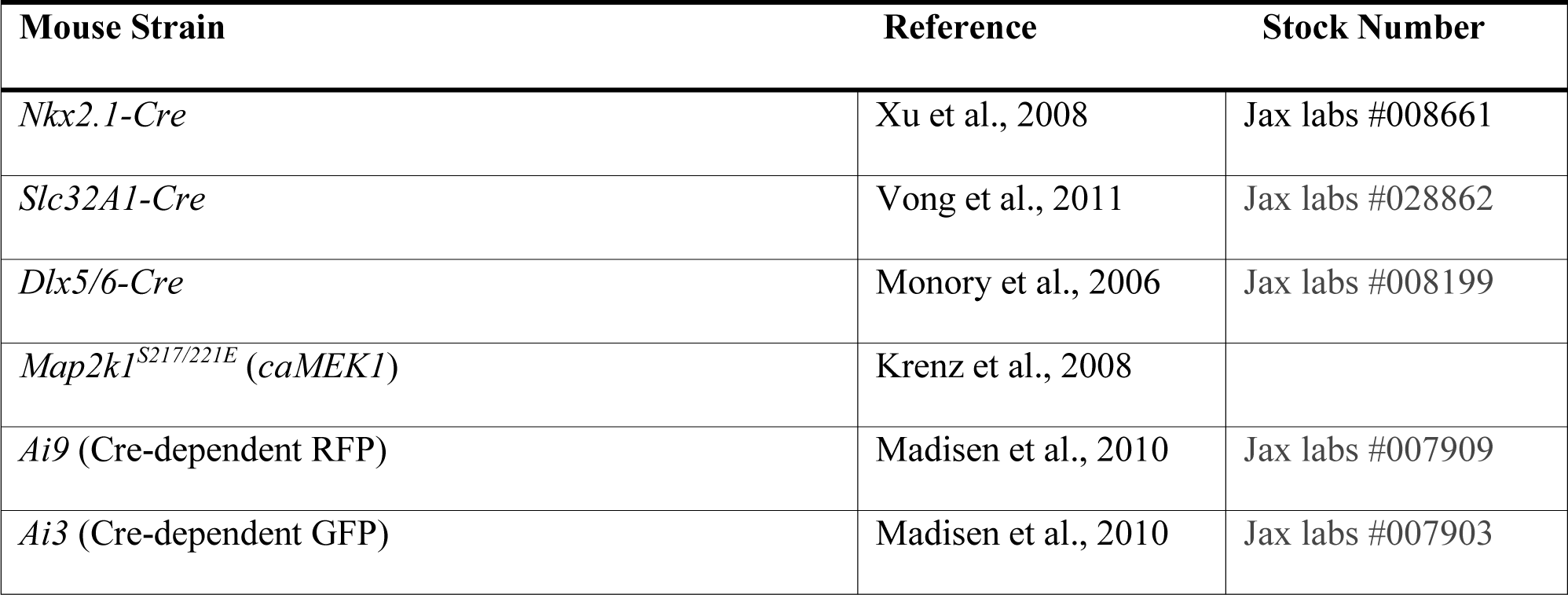
Mouse Strains.

**Table S2.**
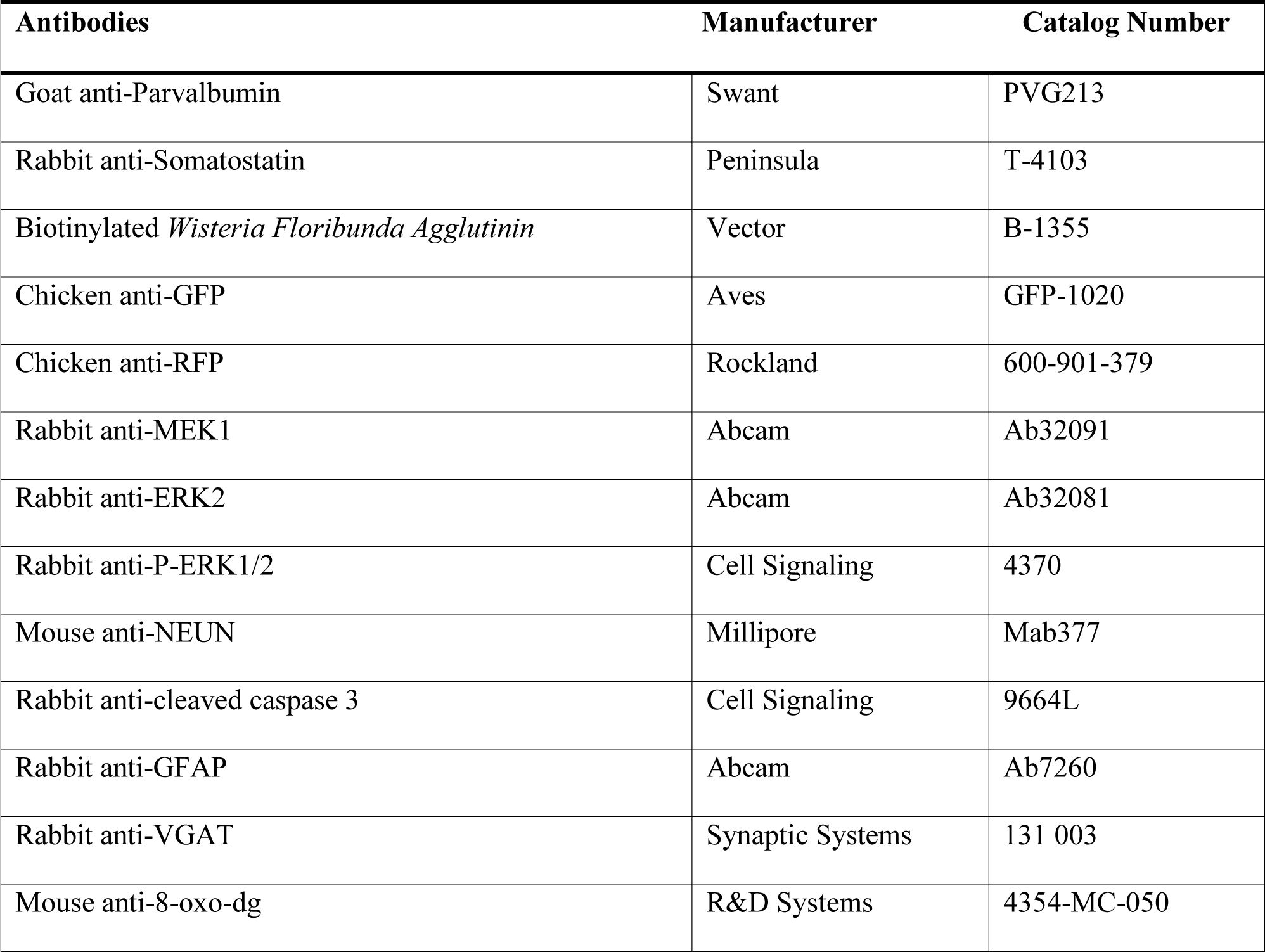
Antibodies.

**Figure S1. Related to Figure 1.**
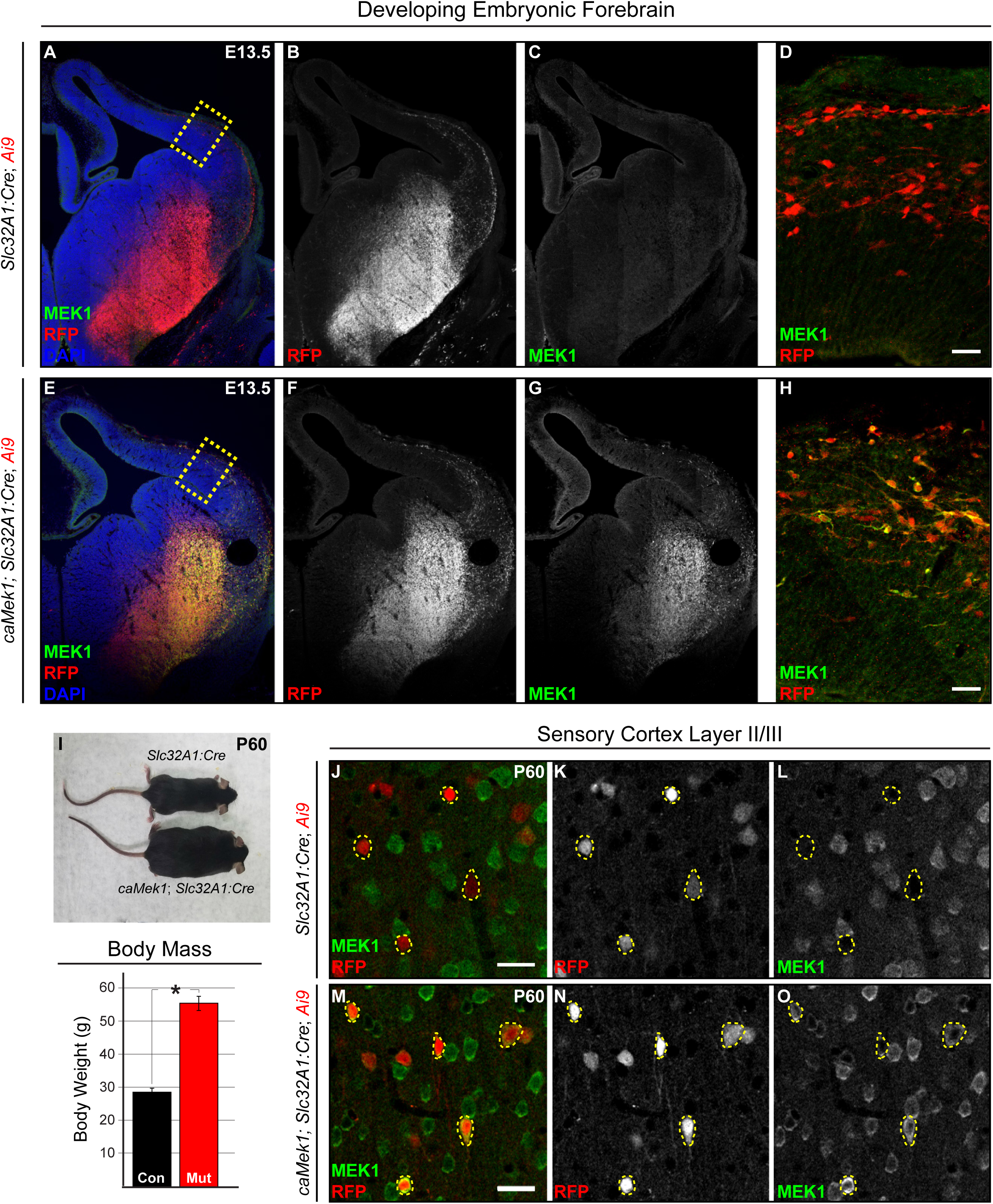
**(A-H)** Representative coronal sections of E13.5 developing mouse forebrain immunolabeled for MEK1. High expression of MEK1 was detected in RFP-labeled cells in *caMek1 Slc32A1:Cre* ganglionic eminences and CIN migratory streams (compare **C** to **G**; **D** to **H**). **(I)** *CaMek1 Slc32A1:Cre* mice exhibited significantly increased body mass in adulthood as compared to controls (n = 12 controls, 12 mutants; mean ± SEM, * = p < 0.05). **(J-O)** Mutant CINs display increased MEK1 expression into adulthood (n=3). (Scale bar = 25 µm)

**Figure S2. Related to Figure 2.**
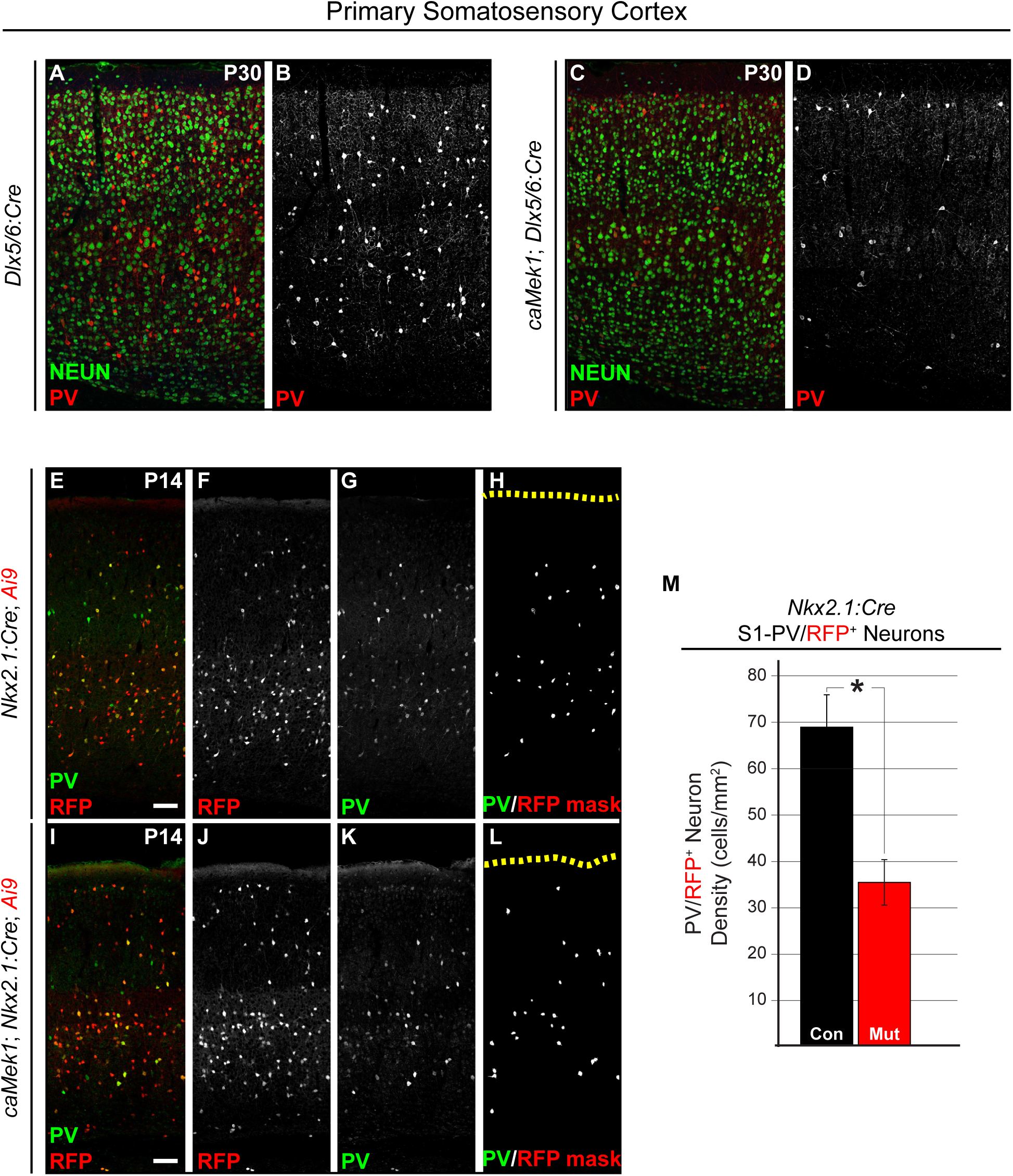
**(A-D)** P30 coronal sections of *caMek1 Dlx5/6:Cre* primary sensory cortex revealed a substantial qualitative decrease in the number of PV-CINs relative to controls (n=3). **(E-M)** Representative confocal images of P14 *caMek1 Nkx2.1:Cre* sensory cortices immunolabeled for PV. The number of PV^+^/RFP^+^ co-expressing cells was significantly decreased in mutants as compared to littermate controls (quantification in **M**: n = 3; mean ± SEM, * = p < 0.05).

**Figure S3. Related to Figure 3.**
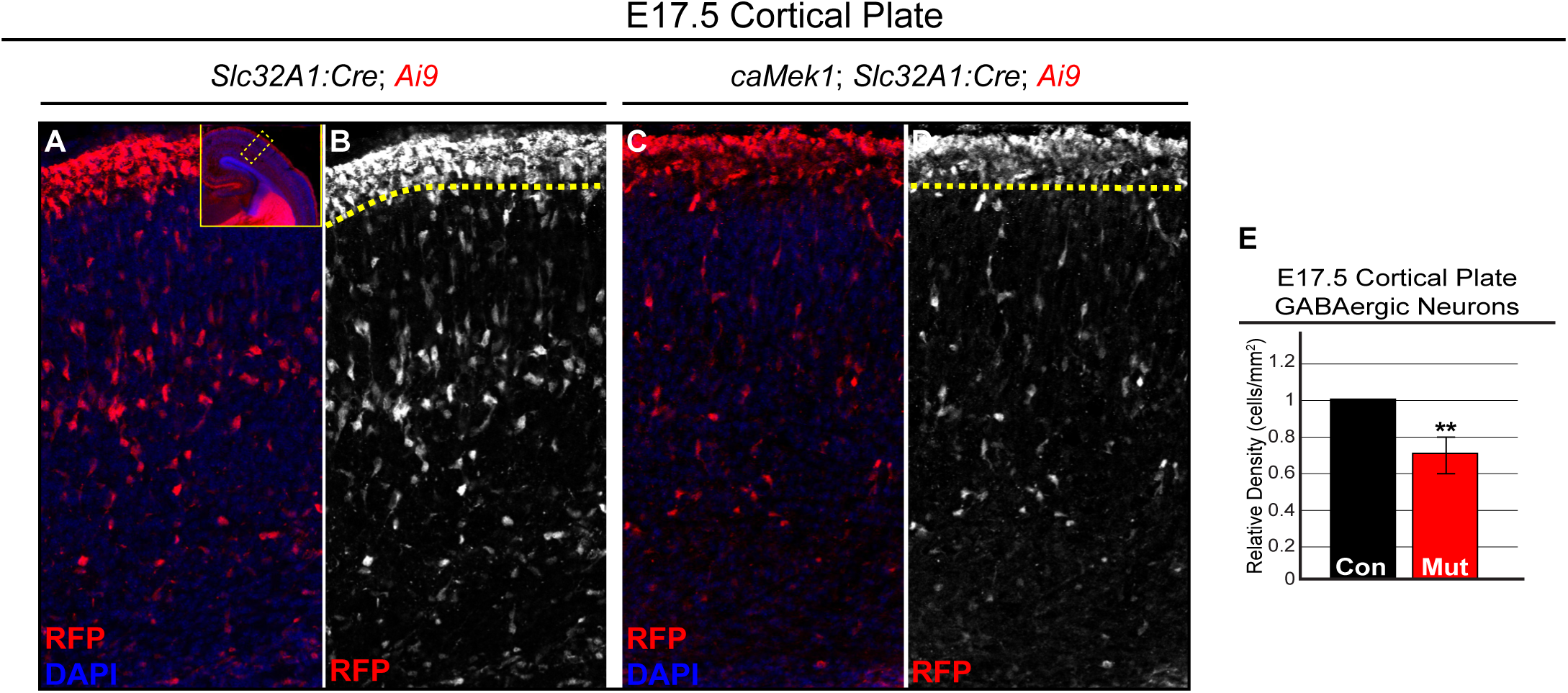
**(A-D)** Representative confocal micrographs of the E17.5 developing cortical plate. A significant decrease in the number of RFP^+^ CINs was detected in *caMek1 Slc32A1:Cre* embryos (quantification in **E**: n = 3; mean ± SEM, * = p < 0.05).

**Figure S4. Related to Figure 4.**
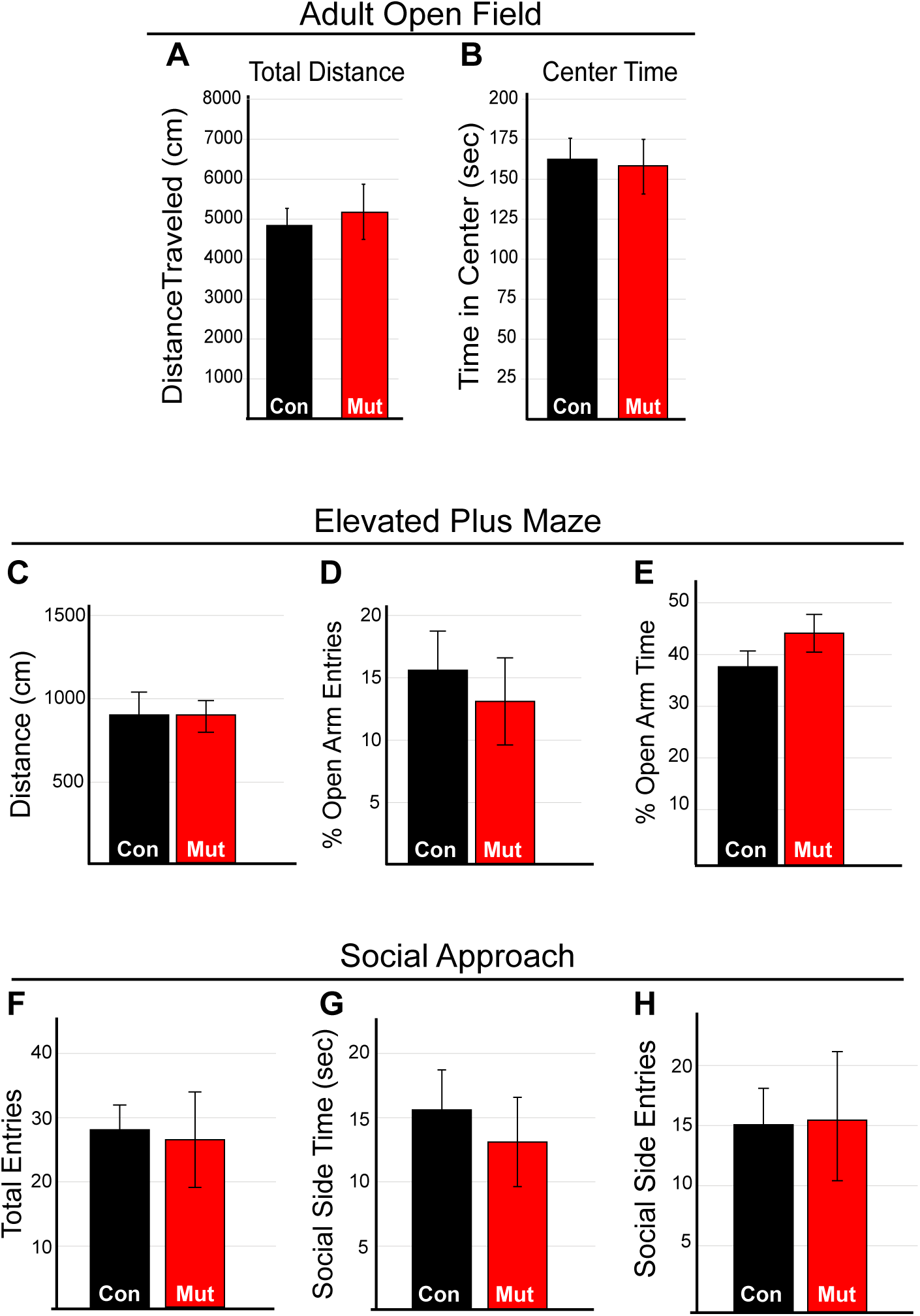
**(A-B)** *CaMek1 Slc32A1:Cre* mice (n = 25 control, 13 mutant) were assessed for locomotor, anxiety-like behaviors, and sociability in the open field task. No significant differences in distance traveled or center time were observed throughout 10 min of open field testing. **(C-E)** Elevated plus maze testing did not detect a significant difference in % open arm entries or % time spent in open arms. **(F-H)** In the social approach assay mutants did not significantly differ from controls in total entries, time spent in the social side, or social side entries.

**Figure S5. Related to Figure 5.**
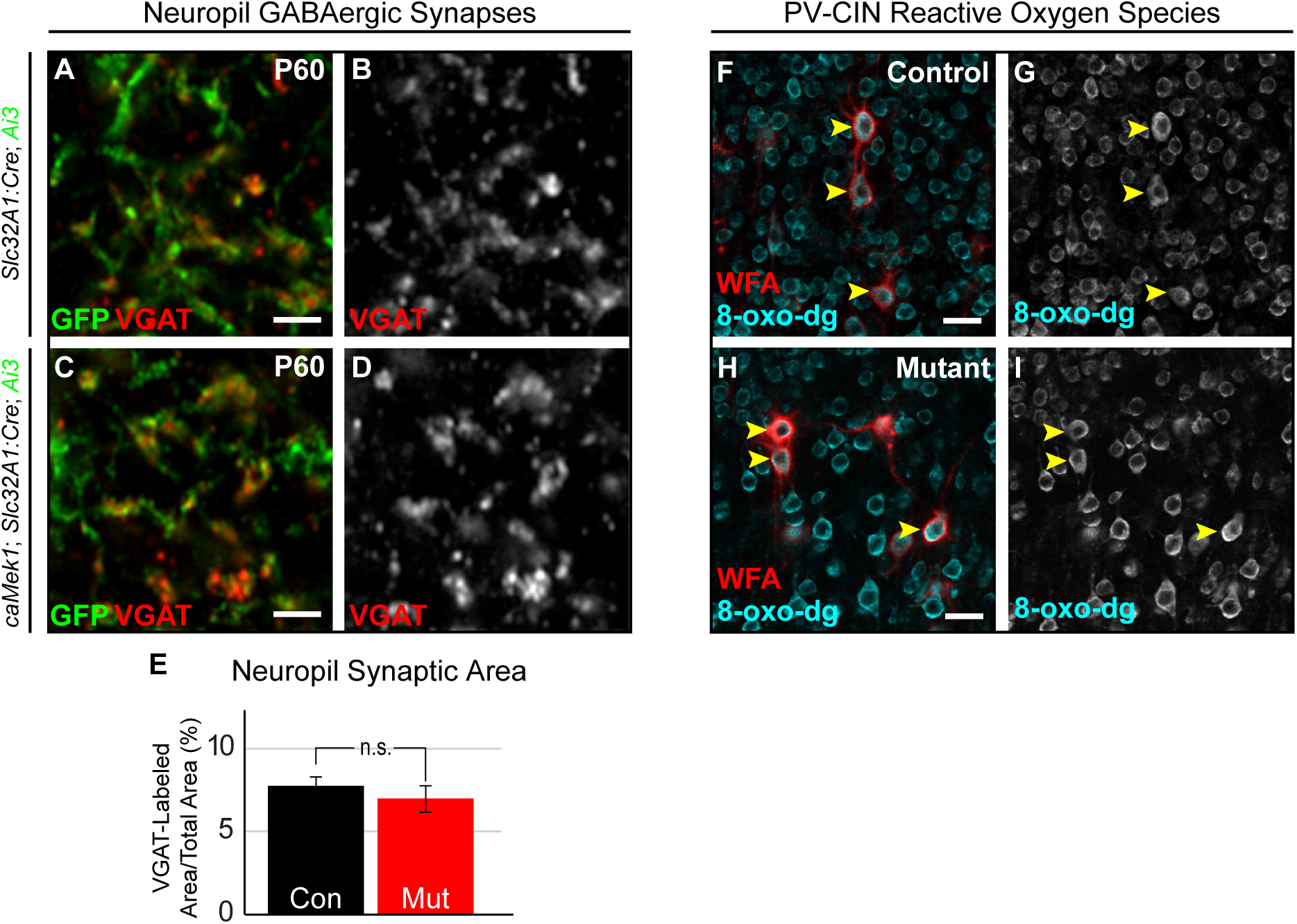
**(A-E)** Mutant mice exhibited normal VGAT-labeling in the layer 2/3 neuropil (quantification in **E**; n = 33 control, 30 mutant regions, mean ± SEM). **(F-I)** Expression of the DNA oxidation marker 8-oxo-dg expression in WFA^+^ *caMek1 Slc32A1:Cre* CINs was qualitatively unchanged when compared to control WFA^+^ CINs (control in F-G, mutant in H-I). (Scale bar = 25 µm).

*Supplemental Video 1.* Representative video montages of two control and two *caMek1 Slc32A1:Cre* mutants that showed abnormal rearing, neck twitching, and hypolocomotion during the first 60 seconds of the open field task.

*Supplemental Video 2.* Representative video montages of three control and three *caMek1 Slc32A1:Cre* mutants that underwent sudden behavioral arrest, abnormal head twitching, or motionless staring during the first 60 seconds of the open field task.

